# Boosting annotation in nutrimetabolomics by Feature-Based Molecular Networking: analytical and computational strategies applied to human urine samples from an untargeted LC-MS/MS based bilberry-blueberry intervention study

**DOI:** 10.1101/2021.12.20.473496

**Authors:** Lapo Renai, Marynka Ulaszewska, Fulvio Mattivi, Riccardo Bartoletti, Massimo Del Bubba, Justin J. J. van der Hooft

## Abstract

Urine represents a challenging metabolite mixture to decipher. Yet, it contains valuable information on dietary intake patterns as typically investigated using randomized, single-blinded, intervention studies. This research demonstrates how the use of Feature-Based Molecular Networking in combination with public spectral libraries, further expanded with an “In-house” library of metabolite spectra, improved the non-trivial annotation of metabolites occurring in human urine samples following bilberry and blueberry intake. Following this approach, 65 berry-related and human endogenous metabolites were annotated, increasing the annotation coverage by 72% compared to conventional annotation approaches. Furthermore, the structures of 15 additional metabolites were hypothesized by spectral analysis. Then, by leveraging the MzMine quantitative information, several molecular families of phase II (e.g., glucuronidated phenolics) and phase I (e.g., phenylpropionic acid and hydroxybenzoic acid molecular scaffolds) metabolism were identified by correlation analysis of postprandial kinetics, and the dietary impact of endogenous and exogenous metabolites following bilberry-blueberry intake was estimated.

## 1. Introduction

Nowadays, untargeted tandem mass spectrometry (MS/MS) is one of the most widely used analytical techniques in metabolomics, allowing for obtaining accurate structural information (i.e., annotation and identification) and occurrence profiles (i.e., quantification) of the chemical compounds present in biological complex mixtures (Schrimpe-Rutledge, Codreanu, Sherrod, & McLean, 2016), thanks also to the coupling with separation techniques such as liquid chromatography (LC). Despite the wide application of hyphenated LC-MS/MS platforms, the annotation of biologically relevant metabolites (i.e., biomarkers) is strongly hampered by the complexity of the metabolome and the decoding procedure applied to this high-dimensional and heterogeneous set of ionized metabolites: i.e., co-eluting and/or not resolved chromatographic peaks of non-isobaric and isobaric compounds, dimers, adducts, in-source fragmentation processes, etc. (Chaleckis, Meister, Zhang, & Wheelock, 2019). Therefore, the annotation process is a pivotal step in untargeted metabolomics, consequently representing a bottleneck for biological information analysis and biomarker discovery. To streamline the metabolite annotation process, standard metabolomics guidelines have been reported for the accurate identification and assignment of a metabolite feature (Kind & Fiehn, 2007; Sumner, Amberg, Barrett, Beale, Beger, Daykin, et al., 2007), through peak picking, mass spectral deconvolution, determination of molecular ions by adduct detection, and fragmentation pattern (MS/MS) analysis (Beniddir, Kang, Genta-Jouve, Huber, Rogers, & van der Hooft, 2021). Furthermore, several tries were undertaken by the scientific community to harmonize not only the annotation steps but also reporting standards: e.g. the MSI (Metabolomics Standards Initiative), FAIR Data Principles (Findable, Accessible, Interoperable and Reproducible), and ELIXIR Metabolomics (Spicer, Salek, & Steinbeck, 2017a, 2017b). Despite these efforts, the risk of missing relevant information and making false discoveries remains relatively high due to incorrect MS and MS/MS interpretations when matching experimental spectra to available spectral libraries. It is also remarkable that the identification of MS/MS similarities within a given dataset is able to highlight the presence of structurally related metabolites, which plausibly share a same metabolic pathway, thus strengthening the biological meaning of the annotations.

In this context, molecular networking (MN) has gained large attention, thanks to the efficient and rapid identification of several molecular families within complex mixtures, providing a visual overview of all the precursor ions of molecules detected and fragmented during an MS experiment grouped according to their structural relationships (Ramos, Evanno, Poupon, Champy, & Beniddir, 2019). MN uses an unsupervised vector-based computational algorithm to organize molecular ions (i.e., clusters or nodes) into a network of molecular families that share spectral similarities among their MS/MS spectra. At the same time, annotation is performed through the Global Natural Products Social Molecular Networking (GNPS) bioinformatics platform (Aron, Gentry, McPhail, Nothias, Nothias-Esposito, Bouslimani, et al., 2020), which represents a unique computational tool linked to a large number of spectral libraries available as public repository of spectra and metadata (i.e., MassIVE), corresponding to putatively annotated (level II) or putatively characterized (level III) compounds, according to metabolomics standards initiative (Sumner, et al., 2007). It is therefore possible to increase the annotation of biologically relevant molecules in comparison with traditional biomarker discovery workflows (Aron, et al., 2020). Indeed, MN has been applied in several untargeted LC-MS/MS studies, mainly focusing on phytochemical composition analysis (Ramos, Evanno, Poupon, Champy, & Beniddir, 2019), and less frequently on drug metabolism (Van Der Hooft, Padmanabhan, Burgess, & Barrett, 2016), and nutrimetabolomics (Said, Truex, Heidorn, Retta, Petrov, Haka, et al., 2020).

Recently, MN has been combined with standard feature detection tools into the Feature-Based Molecular Networking (FBMN) workflow that is capable to resolve isomers and incorporate quantitative information (e.g., spectral counts, chromatographic peak areas, etc.), increasing the link between peak picking algorithms and *in silico* annotation tools (Nothias, Petras, Schmid, Dührkop, Rainer, Sarvepalli, et al., 2020). Until now, FBMN has been successfully applied in various fields of metabolomics, such as the identification of transformation products of organic micropollutants in water samples (Oberleitner, Schmid, Schulz, Bergmann, & Achten, 2021), native plant constituents (Padilla-González, Sadgrove, Ccana-Ccapatinta, Leuner, & Fernandez-Cusimamani, 2020; Rivera-Mondragón, Tuenter, Ortiz, Sakavitsi, Nikou, Halabalaki, et al., 2020; Xie, Kong, Li, Nothias, Melnik, Zhang, et al., 2020) and endogenous urinary metabolites (Neto & Raftery, 2021; Quinn, Melnik, Vrbanac, Fu, Patras, Christy, et al., 2020), while no applications have been reported till now in nutrimetabolomics.

Hence, we hypothesize that the application of state-of-the-art computational metabolomics tools will significantly enhance our understanding of dietary intervention studies by leveraging a larger part of the investigated LC-MS/MS dataset. On this basis, this research aims at investigating the discovery capacity and the possible extension of the annotation coverage of endogenous and exogenous urinary metabolites of the FBMN approach in nutrimetabolomics, in comparison with a conventional identification workflow commonly adopted for discovering significant postprandial metabolites (Renai, Ancillotti, Ulaszewska, Garcia-Aloy, Mattivi, Bartoletti, et al., 2022). The FBMN workflow was applied to deconvoluted and aligned high-resolution LC-MS/MS files of postprandial urine samples from a two-arms intervention study on the intake of *Vaccinium myrtillus* (VM) and *Vaccinium corymbosum* (VC) berry supplements. In detail, samples were analysed in both negative ionization (NI) and positive ionization (PI) data dependent acquisition modes, obtaining high-quality MS/MS spectra, to be compared with the available GNPS libraries. However, MS/MS spectra included in these libraries are obviously obtained under a wide range of instrumental conditions (e.g. time-of-flight, orbitrap, etc.) that may affect the identification process. Accordingly, an “In-house” library containing hundreds of spectra of food-related human metabolites acquired in both PI and NI under the same/similar mass spectrometric conditions of this study, was constructed as elsewhere indicated (Vargas, Weldon, Sikora, Wang, Zhang, Gentry, et al., 2020).

Moreover, thanks to the analysis of quantitative data within networks, a first time attempt to use FBMN for discriminating (i) metabolites characterized by different postprandial kinetics and (ii) the relative dietary contribution of VM and VC interventions in the global occurrence of the identified metabolites, is here presented.

## 2. Materials and methods

### 2.1 Chemicals and reagents

Full purchase detail of solvents and internal and surrogate standards used are reported in section S1 of the *Supplementary materials*. The complete list of the reference standard adopted to build the “In-house” library, in both NI and PI modes, is reported in the *List of Reference Standards used to build the library*.*xlsx* file named in the *Supplementary materials*.

### 2.2 Study design and sample extraction

Urine samples were collected and extracted as previously reported by Ancillotti et al. (Ancillotti, Ulaszewska, Mattivi, & Del Bubba, 2019). Briefly, the randomized, single-blinded two-arms intervention study involved twenty healthy voluntary subjects (11 males and 9 females aged between 25 and 60 years). All the subjects convened early in the morning after 10 h of fasting and were randomly divided in two groups according to an electronic randomisation key. A single dose of 25 g of VM supplement mixed with 500 mL of water was orally administered to the first group, whereas the same amount of VC berry supplement mixed with 500 mL of water was orally administered to the second group. Full details on supplements characterization, i.e. total soluble polyphenols and total monomeric anthocyanins, are reported in section S2 of the *Supplementary materials*. Urine samples of each volunteer were collected at baseline and different sampling times: 30, 60, 120, 240, 360 min and pools at 24h and 48h. Volunteers were asked to avoid fruit and vegetable consumption within the 48 hours after supplement intake.

After the collection, samples were divided in aliquots of 500 µL, frozen at -80°C and stored until extraction and LC-MS/MS analysis were performed. The extraction procedure consisted of filtering and diluting urine samples in accordance with untargeted metabolomics guidelines, and the entire procedure adopted herein is reported in section S3 of the *Supplementary materials*.

### 2.3 LC-MS/MS analysis

LC analysis was performed on a Dionex Ultimate 3000 HPLC system equipped with a Kinetex C18 column (150 mm×2.1 mm I.D., particle size 2.6 μm) and a guard column containing the same stationary phase (Phenomenex, Torrance, CA, USA). The LC system was coupled with a hybrid linear ion trap Fourier Transform (LTQ FT) Orbitrap high-resolution mass spectrometer (Thermo Fisher Scientific, Waltham, MA, USA) by an electrospray ionization (ESI) probe for data-dependent analysis both in positive and negative ionization. Full details of chromatographic and mass spectrometric conditions are reported in section S4 of the *Supplementary materials*.

### 2.4 Metadata

Metadata were entered manually for all samples and organized in two different files according to the polarity adopted. In detail, metadata consisted of three descriptive categories, (i) spectrum file name (the same of acquired raw data), (ii) type of supplement (VM or VC), and (iii) related time point after intake. These elements are required to get a correct grouping within FBMN for quantitative analysis (see the *Metadata and Library Information*.*xlsx* file in the *Supplementary materials*).

### 2.5 Data pre-processing

The data-dependent spectra files (including analysis blanks) were converted from the .raw (Thermo Fisher format) to .mzML open format using MS-Convert software, part of the ProteoWizard package (https://proteowizard.sourceforge.io). Then, to obtain a feature list for NI and PI dataset, the converted files were processed with MZmine 2 software (Pluskal, Castillo, Villar-Briones, & Orešič, 2010) carrying out the following procedures: mass detection for both full scan and MS/MS files, chromatogram building, chromatogram deconvolution, isotope grouping, and alignment. Afterwards, the aligned files are filtered in order to remove the redundant information related to features characterized by similar retention time (Δt_R_ ≤ 0.01 min) and precursor ion mass (Δm ≤ 0.001 Da). Furthermore, the files underwent to the so-called “gap filling procedure”, which has the purpose of making feasible for GNPS to compare acquisitions of different samples, in principle characterized by the presence of signals at different t_R_ and/or *m/z*. In other words, each signal present in at least one sample and absent in at least another, represents for the latter a gap that is filled with a moving average of the baseline signal at that value of *m/z*. Subsequently, the aligned feature list were exported as MS/MS files (.mgf format) and quantification table (.csv format), according to GNPS documentation reported for FBMN workflow (https://ccms-ucsd.github.io/GNPSDocumentation/).

### 2.6 Open format data repository: MassIVE dataset

The study data in mzML format are available on-line on GNPS infrastructure with number MSV000088336. The FBMN analysis are available with following links:

- PI: https://gnps.ucsd.edu/ProteoSAFe/status.jsp?task=a981ebd40809453ebe1524ff1fc8e265
- NI: https://gnps.ucsd.edu/ProteoSAFe/status.jsp?task=0a239e71bb2045c292c4c96f4501249c

### 2.7 “In-house” library implementation

The annotation spread sheets of analytical standards containing their key and machine-readable descriptors such as compound name, SMILES code, InChiKey, PubMed and CAS numbers were used for “In-house” library generation. Two annotation spread sheets were built in NI and PI, containing 319 and 339 injected compounds, respectively (see the *Metadata and Library Information*.*xlsx* file in the *Supplementary materials*). The .mgf file was generated under the GNPS infrastructure for PI and NI modes, after that annotation spread sheets and spectra were validated by launching the “Batch Validator Workflow” (Vargas, et al., 2020). The completed libraries can be found in the public spectral library collection of GNPS under the names “Plant-Human-Nutrition-POS” and “Plant-Human-Nutrition-NEG”.

### 2.8 Molecular networking analyses

Molecular networks were obtained following the online workflow on the GNPS molecular networking web-platform (GNPS) (http://gnps.ucsd.edu). FBMN was performed adopting the most suitable basic and advanced networking options, selected through network optimization by classical MN (see paragraph 3.1), for NI and PI aligned dataset exported from MZmine 2 software. In detail, precursor ion mass tolerance (PIMT) and fragment ion mass tolerance (FIMT) were investigated in the range of 0.01-2 Da and 0.01-1 Da, since they can act as inclusion/exclusion parameters for the clustering of the occurring features and they need to match the dataset well. The number of minimum matched fragment ions to create a consensus was investigated from 2 to 7, as it is also responsible for the creation of a common consensus spectrum. Networking and library cosine scores are parameters related to the similarity degree between consensus and library spectra, used for network creation and annotation, respectively; thus, they were investigated in the range of 0.2-0.7, to test the accuracy of networking and library matching. Finally, the minimum number of library shared peaks was studied from 3 to 5 to reduce the number of incorrect annotations. A comparative analysis looking at library matches, number of molecular families, nodes, and edges, between these settings was performed, tested before and after the inclusion of the “In-house” library, and is reported in **Figure S1** of section S5 of the *Supplementary materials*. The most appropriate input parameters were set as follows: NI were analysed using PIMT = 0.1 Da, FIMT = 0.01 Da, minimum matched fragment ions = 3, networking cosine score > 0.6, library cosine score > 0.5, and minimum library shared peaks = 3. PI dataset was processed adopting PIMT = 0.05, FIMT = 0.05, minimum matched fragment ions = 3, networking cosine score > 0.5, library cosine score > 0.3, and minimum library shared peaks = 3. Blank and quality control (QC) samples were also included in FBMN to check the quality of networks and of quantitative information. Network analysis and quantitative results were investigated and exported adopting Cytoscape environment (Shannon, Markiel, Ozier, Baliga, Wang, Ramage, et al., 2003). Moreover, unknown nodes were annotated with putative molecular structures by manually investigating the following parameters: (i) mass difference between identified and unknown node, (ii) precursor ion mass accuracy, and (iii) fragmentation patterns in MS/MS spectra (see section S5 of *Supplementary materials*).

### 2.9 Analysis of postprandial kinetics

To extract the postprandial information from FBMN, the Pearson product-moment correlation (PPMC) analysis was performed to evaluate the grade of correlation among the chromatographic peak area of identified nodes and time points, using the “corrplot” package implemented in R (https://cran.r-project.org/). The quantitative FBMN data used for the correlation analysis were extracted from the “node table” of the Cytoscape environment, built using the loaded metadata for both the NI and PI datasets. In order to attempt the FBMN related discrimination between phase I and phase II metabolites, the PPMC coefficients (r), which measure the linear dependence between peak area and time points, were used. More in detail, a positive and significant r-value was used to associate nodes to phase I metabolism (i.e., late postprandial metabolites), whereas negative and significant r-values were associated to phase II metabolism (i.e. early postprandial metabolites). For nodes with a not statistically significant coefficient (*P*-value > 0.05), peak area vs. time points plots (data not shown) were manually investigated for a correct postprandial grouping.

To identify the major advantages and/or drawbacks of FBMN, the investigated urine sample underwent traditional biomarker discovery workflow (i.e., statistical inference). Biomarkers of food intake in postprandial responses were selected using full scan data with *R* packages (Garcia□Aloy, Ulaszewska, Franceschi, Estruel□Amades, Weinert, Tor□Roca, et al., 2020), which consisted in a two-step procedure: (i) verification of increasing trend along time points and (ii) calculation of AUC curves and intra-intervention discrimination. For each feature (i.e., a *m/z* at a given retention time), the baseline sample was subtracted from the intensities of the other time points, and negative values were replaced by zero. Features that in at least 2 consecutive points exhibited the 25th percentile of one group higher than the 75th percentile of the other group, were included in the discrimination process based on comparison of area under the curve (AUC). The AUC of each selected feature was calculated between 0 min and 48h time points, using the *pracma* R package, whereas differences of AUCs among diets were tested using the Wilcoxon-Mann Whitney test and the obtained *P*-values were adjusted using the Benjamini Hochberg (BH) method. Adjusted *P*-values□<□0.05 were considered statistically significant.

## 3. Results and discussion

### 3.1 Input parameters for network analysis

Before running FBMN on both NI and PI datasets from VM and VC intervention, various networking basic and advanced options were investigated (i.e. performing MN only) to find out the most suitable parameters to perform the analysis. In order to properly evaluate the effect of input parameters, the total number of nodes, the total number of edges, the number of identified compounds (IDs), i.e., annotated through spectral library matching, and the number of spectral families (i.e., the groups or clusters, also referred to as molecular families), were analysed in both NI and PI datasets. In this regard, increasing the value of precursor ion mass tolerance (PIMT) from 0.01 Da to 2 Da diminished the number of nodes, edges, and spectral families, whereas the number of IDs increased, mainly due to the less strict conditions in merging consensus spectra (i.e., considering different isobaric compounds as one) at increasing PIMT. Hence, to keep a reliable number of nodes without significantly affecting the number of IDs, PIMT was set at 0.1 Da and 0.05 Da for NI and PI, respectively. Fragment ion mass tolerance (FIMT) exerted a similar effect on the output variables as PIMT, apart from the total number of edges, which increased by increasing values of FIMT. Due to the loss of accuracy in nodes networking for high FIMT, analogous values as PIMT were chosen for this parameter. The number of minimum matched fragment ions was set at 3 for both NI and PI, to increase the number of molecular families and/or to make them larger. In this regard, it is also important to note that many food-derived metabolites do only have a few characteristic mass fragments. Cosine scores for networking and library matching were kept below the default value of 0.7, to extract the higher amount of information within NI and PI datasets, which were set at 0.6 and 0.5, respectively. Finally, the number of minimum library shared peaks was set at 3, because higher values of this parameter were responsible of a drastic reduction of IDs, similarly to what observed for the number of matched fragments. **Figs. S1A** and **S1B (**section S5 of the *Supplementary materials*) show the advantageous increase of 20-48% of the identified nodes due to the inclusion of the “In-house” library, and the outputs of final basic and advanced networking conditions, respectively. Subsequently, the selected conditions were adopted for the FBMN workflow. It is worth to mention that, despite the effort to master the matching criteria between fragmentation spectra of library and experimental data, the annotations remained at level II and III, due to the lack of retention time confirmation.

### 3.2 Feature-Based-Molecular-Networking of negative ionisation dataset

FBMN workflow applied to NI dataset through the combination of the aligned feature list and the quantitative tables exported from MZmine 2 software, was able to remove the 57% of redundant IDs (see paragraph 2.5) found by classical MN, which can be a drawback of the latter approach. FBMN of negative ionisation dataset consisted in 545 nodes and 799 edges, with a total number of connected components equal to 307, corresponding to 65 spectral families. **Table 1** reports the 23 identified compounds by library match of nodes within and outside families (i.e., self-loops or singlets). It is remarkable to underline that the 74% of the annotations matched the “In-house” library with high mass accuracy (i.e. mass error < 5 ppm), highlighting the importance of including biochemically relevant metabolites in spectral libraries acquired under experimental conditions comparable with those used for sample analyses. Five “self-loop” nodes were identified, four of them annotated with a good mass accuracy as the following metabolites: azelaic acid, galacturonic acid, glutamine, and ethoxy-oxobutenoic acid. The only exception was the node identified as 13-oxo-7-[3,4,5-trihydroxy-6-(methoxymethyl)oxan-2-yl]oxytetradecanoic acid, which showed a higher mass error (Δ = 87 ppm), thus posing doubts about the goodness of the match. Apart from galacturonic acid and glutamine, which may derive from dissociation phenomena in the ESI source, the other three compounds could be addressed as fatty acid derivatives. The presence of fatty acids in human biofluids has been poorly investigated in association with phenolic-rich fruit interventions. However, azelaic acid and other fatty acids have been detected in human serum after bilberry, pomegranate, and mulberry interventions, probably due to its occurrence in their seeds (Medjakovic & Jungbauer, 2013; Renai, et al., 2022; Zhang, Ma, Luo, & Li, 2018).

**Table 1.** List of the metabolites (ID) annotated by Feature-Based Molecular Networking of negative ionization dataset, including the developed “In-house” library that contains 319 metabolites in negative ionisation mode. Retention time (t_R_, min), mass of precursor ion (MS, Da), type of adduct, product ions (MS/MS, Da) of query spectra and relative base peak in bold, mass error of precursor ion vs. theoretical mass (Δ, ppm), type of library used for annotation, and kind of occurrence in the network (NW), i.e. as singleton (S) or molecular family (MF); n.a. means library match not available.

| # | ID | $t_R$ | MS | Adduct | MS/MS | $\Delta$ | Library | NW |
| --- | --- | --- | --- | --- | --- | --- | --- | --- |
| 1 | Hydroxyphenyl propionic acid | 4.95 | 165.056 | M-H | **121.066**; 121.028 | 0.5 | “In-house” | MF |
| 2 | Dihydroxyphenyl propanoic acid glucuronide | 3.10 | 357.083 | M-H | 341.272; 175.0248; **113.0245** | 2.9 | “In-house” | MF |
| 3 | Methoxyindole acetic acid | 6.40 | 204.067 | M-H | **160.077**; 130.066; 117.071 | 1.6 | “In-house” | MF |
| 4 | Hydroxy-methoxycinnamic acid | 4.80 | 193.051 | M-H | **178.027**; 149.061; 134.037 | 0.2 | “In-house” | MF |
| 5 | Isovanillic acid | 3.71 | 167.035 | M-H | 152.011; **123.045**; 108.022 | 0.6 | “In-house” | MF |
| 6 | α-Hydroxyhippuric acid | 4.73 | 194.046 | M-H | **150.056** | 1.03 | “In-house” | MF |
| 7 | Azelaic acid | 6.54 | 187.098 | M-H | **125.0972** | 1.5 | “In-house” | S |
| 8 | Dihydrocaffeic acid sulphate | 4.44 | 261.007 | M-H | **181.050** | 1.8 | “In-house” | MF |
| 9 | Dihydroferulic acid sulphate | 4.66 | 275.023 | M-H | **195.066** | 2.6 | “In-house” | MF |
| 10 | Furoylglycine | 2.61/2.91 | 168.03 | M-H | **124.040**; 100.004; 67.019 | 2.2 | “In-house” | MF |
| 11 | Galacturonic acid | 6.38 | 193.035 | M-H | 199.888; 175.025; 131.035; **113.025** | 2.5 | “In-house” | S |
| 12 | Glutamine | 4.81 | 145.062 | M-H | 128.036; **127.052**; 109.041 | 1.4 | “In-house” | S |
| 13 | Hippuric acid | 4.74 | 178.051 | M-H | **134.0609** | 0.9 | “In-house” | MF |
| 14 | Homovanillic acid sulphate | 4.00 | 261.007 | M-H | 217.017; **181.050**; 137.061 | 1.8 | “In-house” | MF |
| 15 | Indoxyl sulphate | 4.75 | 212.002 | M-H | 132.045; **80.965**; 79.958 | 0.9 | “In-house” | MF |
| 16 | Isoferulic acid glucuronide | 4.92 | 369.083 | M-H | 193.050; 175.024; **113.024**; 95.0139 | 1.7 | “In-house” | MF |
| 17 | Dihydrocaffeic acid glucuronide | 3.90/4.10 | 357.082 | M-H | **313.093**; 175.025; 113.025 | 1.7 | “In-house” | MF |
| 18 | Ethoxy-oxobutenoic acid | 2.87 | 143.035 | M-H | 99.045; 81.035; **71.014** | 0.7 | Massbank | S |
| 19 | Enterolactone | 6.79 | 297.113 | M-H | **253.123**; 189.0552; 165.056;<br>121.066 | 0.3 | Massbank | MF |
| 20 | Dihydroxybenzene sulphate | 3.73 | 188.986 | M-H | **109.030**; 80.965; 79.958 | 0.6 | Massbank | MF |
| 21 | Methoxyabscisic acid glucuronide | 7.09 | 453.175 | M-H | 435.166; **277.144**; 233.155; 175.025 | 2.6 | n.a. | MF |
| 22 | 13-oxo-7-[3,4,5-trihydroxy-6-(methoxymethyl)oxan-2-yl]oxytetradecanoic acid | 7.60 | 433.245 | M-H | 415.197; **257.175**; 239.1650 | 87 | Dorrestein | S |
| 23 | p-Cresol sulphate | 5.64 | 187.009 | M-H | 187.007; 107.057; **79.958** | 1.1 | A.Walker | MF |

The other 18 annotated compounds occurred inside molecular families, allowing for identifying relevant metabolite families. **Figures 1** and **2** show the spectral families in which was annotated at least one metabolite, and in some cases unknown nodes with hypothesized compound structures. Moreover, the quantitative information about the contribute of VM and VC interventions (i.e., spectral counts contributing to each node) is reported with a pie chart in each node, whereas postprandial kinetics is highlighted by red and blue boxes and by the reported statistically significant correlation coefficients. In detail, **Fig. 1** reports spectral families belonging to the category of phenolic acids glycosides, most of them identified as glucuronide metabolites, thanks to the occurrence in MS/MS spectra of peaks at *m/z* 175.02 and 113.02, which are typical of glucuronic acid. The principal molecular scaffolds occurring in 9 out of 11 nodes were related to cinnamic and dihydrocinnamic acids, i.e. dihydroxyphenyl propanoic acid glucuronide, isoferulic acid glucuronide, dihydrocaffeic acid glucuronide, hydroxyphenyl propionic acid, and enterolactone. **Figure S2** of the *Supplementary materials* shows the MS/MS spectra of the unknown nodes inside **Fig. 1** molecular families (labelled with a gear), which allowed to putatively identify structurally similar metabolites belonging to these compound categories, as pointed out by the hypothesized scheme of fragmentation (**Fig. S2**). The remaining two nodes were recognized as a molecular family related to abscisic acid glucuronide derivatives (**Fig. 1**). The identified node present in this family, was at first addressed as dihydroxy-diphenylphenoxy-trihydroxyoxane-carboxylic acid, with a mass error of about 128 ppm. The manual investigation of its MS/MS spectra (see **Figure S3** of the *Supplementary materials*) led to a more accurate putative annotation of this node as methoxyabscisic acid glucuronide (Δ = 2.6 ppm). In addition, the hypothesized structure of the linked node was consistent with abscisic acid glucuronide (**Fig. S2**), which was already putatively identified in a previous study (Ancillotti, Ulaszewska, Mattivi, & Del Bubba, 2019) by traditional annotation workflows. The analysis of postprandial kinetics of the spectral families reported in **Fig. 1** allowed for highlighting a phase II expression (r = −0.302) related to dihydroxyphenyl propanoic acid glucuronide, thus suggesting its direct origin from the supplement (Ancillotti, Ciofi, Rossini, Chiuminatto, Stahl-Zeng, Orlandini, et al., 2017). Differently, a mixed contribution of phase I and phase II metabolisms can be suggested for the other identified molecular families. In detail, as regards isoferulic acid glucuronide, even though it shows a positive and significant PPMC correlation (r = 0.757), it is linked with a node exhibiting an opposite postprandial behaviour (peak area vs. time points, data not shown), thus suggesting a phase I-II mixed contribution. An analogous mixed metabolic contribution can be proposed for the abscisic acid spectral family since the methoxyabscisic acid glucuronide is most likely associated to phase I due to the methylation of the hydroxyl group, while the node manually investigated and putatively associated to the glucuronide derivative of the plant-native abscisic acid, is related to phase II. The spectral family associated with the phenylpropionic scaffold also included metabolites associated with both phase I (i.e., enterolactone and hydroxyphenylpropionic acid) and phase II (i.e., dihydrocaffeic acid glucuronide) (Vetrani, Rivellese, Annuzzi, Adiels, Borén, Mattila, et al., 2016). **Fig. 2A** shows the spectral family of several sulfate metabolites belonging to the categories of dihydrocinnamic and vanillic acids, phenolic derivatives, and indoles. Thanks to the analysis of postprandial profiles and in agreement with literature findings (Aravind, Wichienchot, Tsao, Ramakrishnan, & Chakkaravarthi, 2021; Stevens & Maier, 2016), the molecular scaffolds of the identified compounds are probably related to the activity of the gut microbiota. Hence, the metabolites occurring in this spectral family can be addressed to the phase I metabolism (0.526 < r < 0.700). **Fig. 2B** illustrates the spectral family of α-hydroxyhippuric acid, hydroxyphenyl propionic acid, hippuric acid, furoylglycine, hydroxy-methoxy cinnamic acid, methoxyindole acetic acid, and isovanillic acid. Interestingly, apart from the first three listed compounds, which are well-known biomarkers of VM intake (Ancillotti, Ulaszewska, Mattivi, & Del Bubba, 2019; de Mello, Lankinen, Lindström, Puupponen□Pimiä, Laaksonen, Pihlajamäki, et al., 2017) and generic markers of polyphenol-rich dietary intake, the metabolites shown in **Fig. 2B** represent the first report associated with human consumption of bilberry and blueberry. Moreover, the postprandial analysis of the identified nodes evidenced a phase I expression (0.348 < r < 0.940) for most metabolites, with the only exception of hydroxyphenyl propionic acid, for which a phase II metabolism can be suggested, based on its r-value (−0.415). Even though this metabolic association is apparently questionable due the lack of a conjugated group, the analysis of the full scan acquisition evidenced an in-source fragmentation of the sulfate derivative of the hydroxyphenyl propionic acid (*m/z* = 263.02 Da), thus confirming the phase II metabolic attribution.

**Figure 1.**
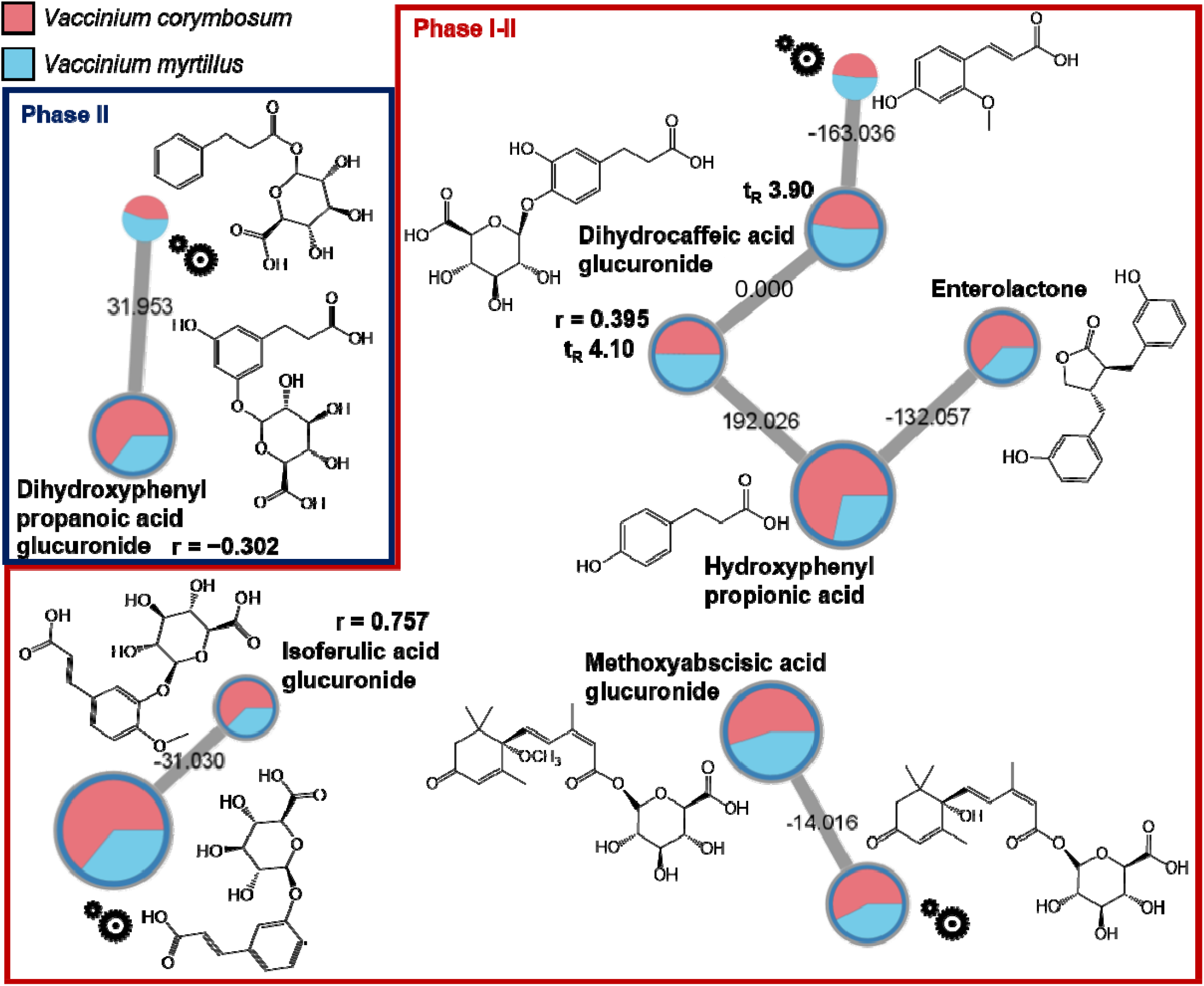
Extracted networks of identified metabolites in negative ionisation mode listed in Table 1, belonging to the category of (poly)phenolic compounds, abscisic acid, and their glucuronide derivatives. Blue and red boxes highlight the metabolism of phase II and I-II, respectively. The “gear” symbols refer to the putative structure identified by manual investigation as reported in paragraph 2.8 of the main manuscript. Statistically significant correlation coefficients (r) are reported. Edge labels refer to the mass difference between two nodes.

**Figure 2.**
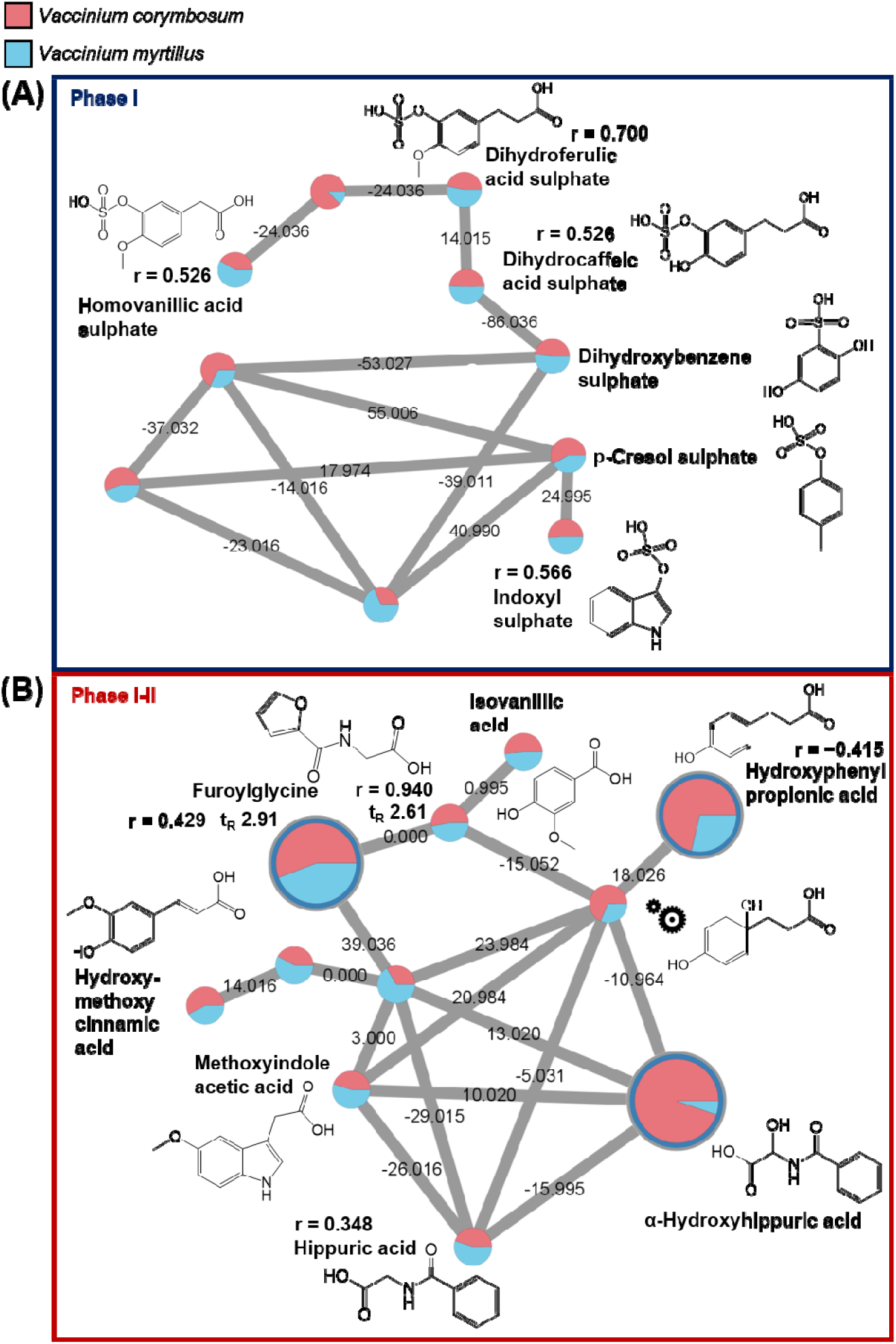
Extracted network of identified metabolites in negative ionisation mode, belonging to the category of (A) sulfate, (B) (poly)phenolic and general metabolites. Blue and red boxes highlight the metabolism of phase II and I-II, respectively. The “gear” symbol refers to the putative structure identified by manual investigation as reported in paragraph 2.8 of the main manuscript. Statistically significant correlation coefficients (r) are reported. Edge labels refer to the mass difference between two nodes.

The structure of one unknown node was also hypothesized in this family (phenylpropionic scaffold), thanks to the resulting mass difference against hydroxyphenyl propionic acid (**Fig. 2B**) and the analysis of MS/MS fragmentation pattern reported in **Figure S4** of the *Supplementary materials*. the quantitative information about the contribute of VM and VC interventions (i.e., spectral counts contributing to each node) is reported with a pie chart in each node

Finally, by summing IDs spectral counts (i.e., the number of spectra with the same consensus spectrum) contributing to each VM and VC intervention, it was possible to evaluate the general impact on the global urine metabolome. Overall, it resulted in a 62% of IDs postprandial occurrence after the intake of VC supplement, and in a 38% for VM.

### 3.3 Feature-Based-Molecular-Networking of positive ionisation dataset

FBMN workflow applied to PI dataset was able to remove 27% of redundant IDs (see paragraph 2.5) found by classical MN mostly likely by removing artefacts like duplicated features. FBMN of PI dataset consisted in 5079 nodes and 6904 edges, with a total number of connected components equal to 3543 (i.e., 663 spectral families). **Table 2** reports the 42 identified compounds by library match among molecular families and singlets. Unlike in the NI dataset, the percentage of annotated compounds by “In-house” library was equal to 43%, due to the greater number of available libraries in PI mode with a mass accuracy of below 5 ppm for most spectral matches.

**Table 2.** List of the metabolites (ID) annotated by Feature-Based Molecular Networking of positive ionization dataset, including the developed “In-house” library, that contains 339 metabolites in positive ionisation mode. Retention time (t_R_, min), mass of precursor ion (MS, Da), type of adduct, product ions (MS/MS, Da) of query spectra and relative base peak in bold, mass error of precursor ion vs. theoretical mass (Δ, ppm), type of library used for annotation, and kind of occurrence in the network (NW), i.e. as singleton (S) or molecular family (MF).

| # | ID | $t_R$ | MS | Adduct | MS/MS | $\Delta$ | Library | NW |
| --- | --- | --- | --- | --- | --- | --- | --- | --- |
| 1 | Tetrahydroharmane carboxylic acid | 4.79 | 231.113 | M+H | **214.086**; 188.070; 158.096 | 0.9 | "In-house" | MF |
| 2 | Hydroxy-dimethoxyphenyl-ethanone | 5.60 | 197.080 | M+H | 169.086; **151.075**; 133.101 | 5.0 | BMDMS-NP | MF |
| 3 | 1,6-propano-ethylideno-1,4-piperazine-2,5-dione | 2.04 | 181.096 | M+H | 153.102; 149.023; **125.071** | 4.5 | Liaw-Yang | MF |
| 4 | Dimethyl-uric acid | 3.51 | 197.067 | M+H | 182.043; 158.056; **140.045** | 5.1 | Dorrestein | MF |
| 5 | β-Glucopyranosyl-tryptophan | 1.90 | 367.150 | M+H | 349.139; 331.129; **247.108**;<br>229.097 | 0.8 | C. Maier | S |
| 6 | Dihydroxycinnamic-trihydroxycyclohexane carboxylic acid | 4.54 | 355.102 | M+H | 341.216; **285.010**; 266.999;<br>163.039 | 4.7 | MoNA | S |
| 7 | Tetrahydro-β-carboline carboxylic acid | 4.52 | 217.097 | M+H | 173.107; 171.091; **144.081** | 0.2 | "In-house" | MF |
| 8 | Dihydroxyphenylglycol | 6.06 | 153.054 | M-H<sub>2</sub>O+H | 135.116; **107.049** | 2.0 | Dorrestein | MF |
| 9 | Methoxyphenyl-propionic acid | 8.13 | 163.075 | M-H<sub>2</sub>O+H | 135.080; 121.1006; **107.085** | 2.4 | "In-house" | MF |
| 10 | Methoxy-hydroxymandelate | 2.03 | 181.049 | M-H<sub>2</sub>O+H | 153.055; **149.023**; 125.071 | 5.0 | Dorrestein | MF |
| 11 | Ferulic acid | 5.14 | 177.054 | M-H<sub>2</sub>O+H | **145.028** | 3.0 | "In-house" | MF |
| 12 | Vanillic acid | 3.65 | 169.050 | M+H | **125.059**; 93.033 | 0.5 | "In-house" | MF |
| 13 | Dihydroxy-trimethyl-isochromenone | 7.37 | 223.096 | M+H | 205.085; **181.086**; 163.075 | 2.3 | MoNA | S |
| 14 | Absicic Acid | 7.38 | 247.133 | M-H<sub>2</sub>O | **229.122**; 211.111; 201.127;<br>187.111 | 0.4 | "In-house" | S |
| 15 | Alpha-CEHC | 6.96 | 261.148 | M-H<sub>2</sub>O+H | **243.138**; 165.091 | 1.6 | "In-house" | S |
| 16 | Azelaic acid | 6.55 | 171.101 | M-H<sub>2</sub>O+H | 153.090; **125.096** | 2.2 | "In-house" | S |
| 17 | Cinnamic acid | 4.86 | 149.060 | M+H | **131.049** | 4.0 | BMDMS-NP | S |
| 18 | Curcumenol | 6.86 | 235.169 | M+H | **217.158**; 189.163; 137.0592;<br>111.0800 | 0.4 | MoNA | MF |
| 19 | Dihydro-deoxy-lactucin | 6.41 | 263.129 | M+H | **245.117**; 217.122; 188.091 | 6.5 | HTSimonsen | S |
| 20 | Ethylindole carboxylic acid | 8.01 | 190.086 | M+H | 172.075; **144.080**; 118.065 | 0.2 | "In-house" | S |
| 21 | Folinic acid | 3.78 | 474.173 | M+H | 456.162; **345.130**; 327.119 | 1.5 | "In-house" | S |
| 22 | Formylkynurenine | 2.14 | 237.087 | M+H | **220.061**; 202.050; 192.066 | 3.8 | Quinn | S |
| 23 | Furaneol | 4.03 | 129.055 | M+H | 111.044; **101.059** | 2.5 | "In-house" | S |
| 24 | Furoylglycine | 2.68 | 152.034 | M-H<sub>2</sub>O+H | **124.039**; 95.013 | 3.5 | "In-house" | S |
| 25 | Gamma-CEHC | 6.68 | 247.132 | M-H<sub>2</sub>O | **229.122**; 203.106; 151.075 | 2.7 | "In-house" | MF |
| 26 | Indole acetic acid | 5.33 | 176.071 | M+H | 145.028; **131.068** | 2.3 | M. Ernst | S |
| 27 | Isoferulic acid | 4.91 | 177.054 | M-H<sub>2</sub>O+H | 163.039; **145.028**; 117.033 | 3.0 | "In-house" | MF |
| 28 | Methylcinnamate | 4.96 | 163.075 | M+H | **131.075**; 103.054 | 1.2 | Massbank | MF |

**Table 2** — Continued
| # | ID | t<sub>R</sub> | MS | Adduct | MS/MS | Δ | Library | NW |
| --- | --- | --- | --- | --- | --- | --- | --- | --- |
| 29 | Caffeine | 4.63 | 195.088 | M+H | **138.066** | 6.1 | Massbank | MF |
| 30 | Trihydroxybutyrophenone | 6.37 | 197.081 | M+H | 169.086; **151.075**; 137.059 | 1.5 | Massbank | S |
| 31 | Hesperetin | 6.70 | 303.086 | M+H | 285.186; **177.054**; 153.018 | 0.7 | Massbank | S |
| 32 | Dihydroresveratrol | 6.19 | 231.102 | M+H | **213.148**; 203.1062; 137.059;<br>121.064 | 3.5 | Massbank | S |
| 33 | m-Coumaric acid | 5.21 | 147.044 | M-H<sub>2</sub>O+H | **119.049**; 105.069 | 1.3 | "In-house" | MF |
| 34 | Dihydroxy-trimethyl-dihydro-isochromen-one | 4.29 | 223.096 | M+H | **205.086**; 163.075 | 4.0 | Dorrestein | MF |
| 35 | Nerol | 8.26 | 137.132 | M-H<sub>2</sub>O+H | **95.085**; 81.070 | 2.7 | "In-house" | S |
| 36 | p-Cumaric acid | 4.04 | 147.044 | M-H<sub>2</sub>O+H | 130.050; **119.049** | 2.0 | "In-house" | MF |
| 37 | Sebacic acid | 7.33 | 185.117 | M-H<sub>2</sub>O+H | **167.106**; 121.101 | 2.9 | "In-house" | S |
| 38 | Ketodeoxycholic acid | 8.14 | 373.274 | M+H-H<sub>2</sub>O | **355.262**; 337.252 | 1.9 | R. Knight | S |
| 39 | Keto-octadecadienoic acid | 8.05 | 295.226 | M+H | **277.216**; 249.221 | 0.9 | R. Knight | S |
| 40 | Hydroxy-methoxybenzophenone | 7.17 | 229.087 | M+H | **151.039**; 105.033 | 5.3 | W. H.Gerwick | S |
| 41 | Enterolactone | 6.78 | 281.117 | M+H-H<sub>2</sub>O | **263.106**; 149.060; 133.065 | 2.8 | R. Knight | S |
| 42 | Ligustilide | 7.59 | 191.107 | M+H | 163.075; **145.101**; 117.070 | 2.6 | J. L.<br>Wolfender | MF |

Twenty-four singlets were identified and, interestingly, they belong to very different categories of endogenous and exogenous metabolites. In detail, by manually plotting features peak area vs. time points, several (poly)phenolics and phenolics derivatives were annotated in association with phase I metabolism (i.e. dihydroxy-trimethyl-isochromenone, trihydroxybutyrophenone, and dihydroresveratrol), phase II metabolism (i.e. dihydroxycinnamic-trihydroxycyclohexane carboxylic acid), and mixed contribution of phase I-II (i.e. cinnamic acid and hesperetin). Additional metabolic contributions were highlighted by plant endogenous compounds, such as β-glucopyranosyl-tryptophan, furaneol, which are already known as food-intake biomarker (0.201 < r < 0.413) (Du, Finn, & Qian, 2010; Gutsche, Grun, Scheutzow, & Herderich, 1999), and abscisic acid and nerol as plant constituents (Degu, Ayenew, Cramer, & Fait, 2016; Elsharif & Buettner, 2016), hence confirming their observed early occurrence in urine samples (−0.214 < r < −0.275). The 9 identified human endogenous compounds (i.e., alpha-CEHC, ethylindole carboxylicacid, folinic acid, formylkynurenine, indole acetic acid, sebacic acid, ketodeoxycholic acid, keto-octadecadienoic acid, and hydroxy-methoxybenzophenone) exhibited a varied correlation trend against time points (−0.645 < r < 0.958), resulting in a complex metabolic output in association with the investigated interventions. Finally, PI mode exhibited three IDs that matched the NI annotations of azelaic acid, furoylglycine and enterolactone, characterized by high correlation coefficients (0.640 < r < 0.967). The other annotated compounds occurred inside molecular families (**Table 2**), allowing for identifying interesting metabolites. **Figures 3** and **4** display the spectral families of metabolites occurring in PI dataset. On the left side of **Fig. 3**, two nodes were identified as isomers of vanillic acid at different retention times, thanks to the spectral match with the “In-house” library. In addition, the remaining nodes were putatively addressed as vanillic acid derivatives with high mass accuracy (from –3.26 to –0.06 ppm), the structures of which were hypothesized by the analysis of MS/MS spectra (see **Figure S5** of the *Supplementary materials*). This molecular family has been associated to protocatechuic acid derivatives, which resulted to belong to both phase II and I metabolisms. In fact, vanillic acid was characterized by a negative correlation against time points (i.e., r = −0.197), suggesting its direct origin from the supplements intake, as also confirmed by literature findings (Colak, Primetta, Riihinen, Jaakola, Grúz, Strnad, et al., 2017). On the other hand, the two hypothesized protocatechuic acid derivatives exhibited an increasing signal around 6-24 hours (data not shown), probably due to the interaction in the large intestines of microbiota with supplement native polyphenols (i.e., anthocyanins and flavonols), resulting in their breakdown to small phenolic acids (Stevens & Maier, 2016). On the right side of **Fig. 3**, the three nodes occurring in the spectral family were identified as dihydroxy-trimethyl-isochromenone and two isomers of ligustilide. The first metabolite was characterized by a negative and significant correlation coefficient (r = −0.164) and an in-source fragmentation of its sulfate derivative was observed by analysing the full scan acquisition, thus suggesting a phase II metabolic origin. Conversely, ligustilide exhibited an opposite correlation trend (r = 0.531) compared to isochromenone, and could be therefore associated to a phase I metabolism. In this regard, even though the two metabolites were included in the same spectral family, no metabolic and/or structural relation seem to occur between them, thus posing some doubts about their FBMN library matching.

**Figure 3.**
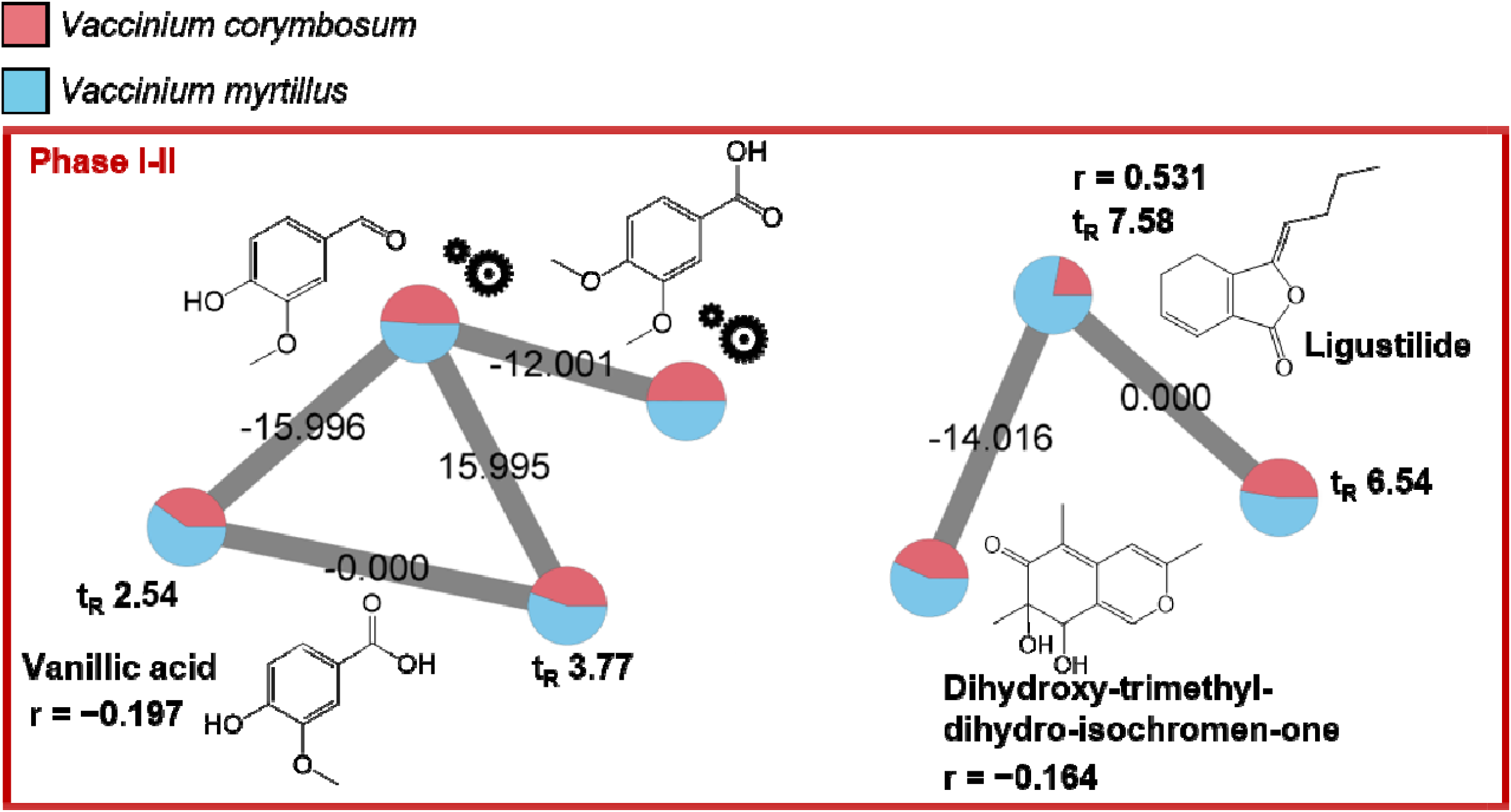
Extracted networks of identified metabolites in positive ionisation mode listed in Table 2, belonging to the category of (poly)phenolic derivatives and plant endogenous constituents. The “gear” symbols refer to the putative structure identified by manual investigation as reported in paragraph 2.8 of the main manuscript. Statistically significant correlation coefficients (r) are reported. Edge labels refer to the mass difference between two nodes.

**Figure 4.**
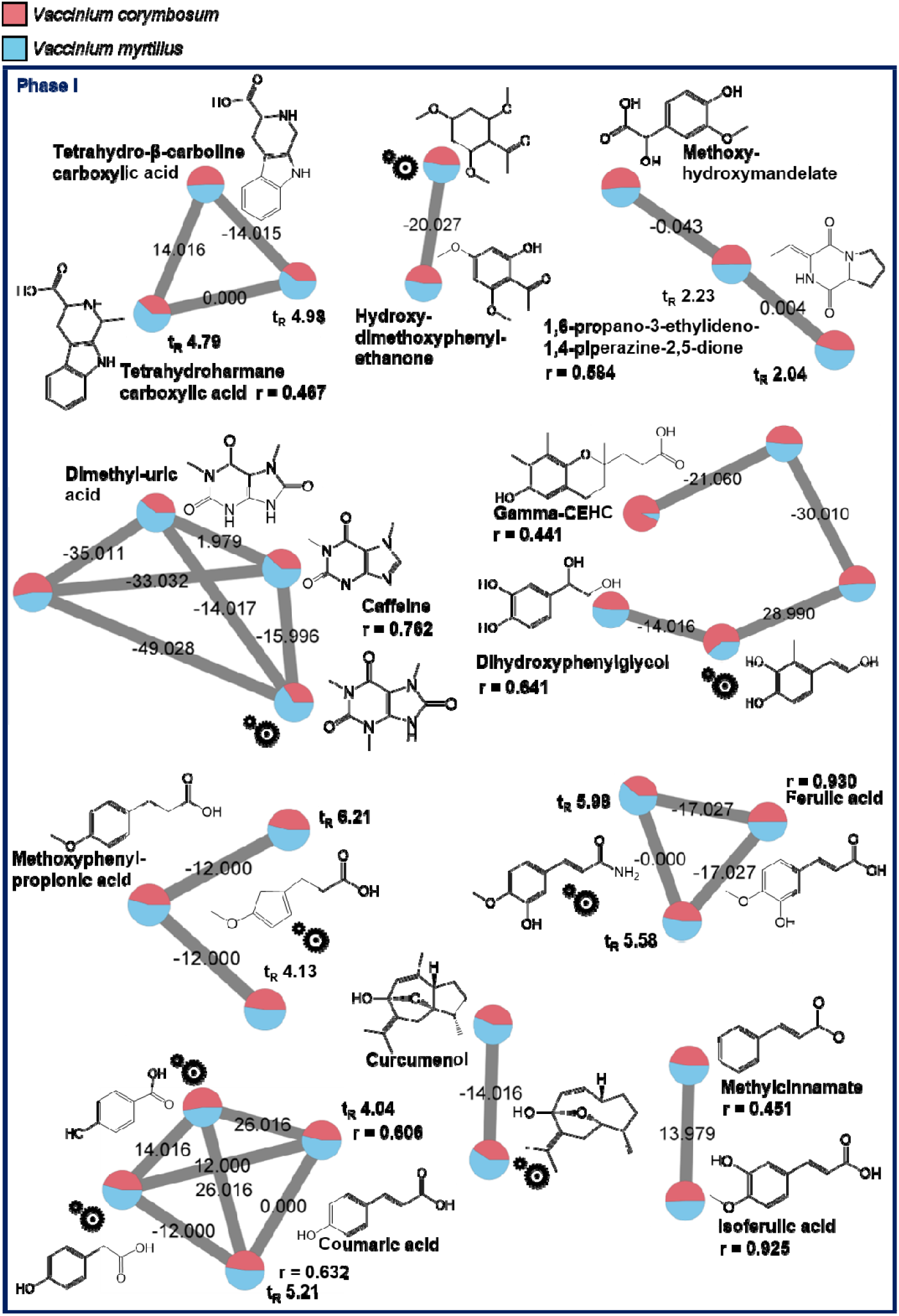
Extracted networks of identified metabolites in positive ionisation mode listed in Table 2, belonging to the category of (poly)phenolic and plant endogenous constituent derivatives occurring in the phase I metabolism. The “gear” symbols refer to the putative structure identified by manual investigation as reported in paragraph 2.8 of the main manuscript. Statistically significant correlation coefficients (r) are reported. Edge labels refer to the mass difference between two nodes.

**Fig. 4** reports a wide number of spectral families, which could be related to phase I metabolism based on postprandial kinetics analysis (0.441 < r < 0.930). The 42% of the identified and hypothesized node structures can be addressed as phase I metabolite of the native polyphenols occurring in the bilberry and blueberry supplements (Stevens & Maier, 2016). In detail, four families can be fully recognized as phenolic metabolites, namely derivatives of phloroglucinol carboxylic acid (i.e., hydroxy-dimethoxyphenyl-ethanone), cinnamic acid (i.e., coumaric acid, methylcinnamate, ferulic and isoferulic acid), and mandelic acid (i.e., methoxy-hydroxymandelate). Moreover, three interesting spectral families were identified: β-carboline derivatives (i.e., tetrahydroharmane carboxylic acid and tetrahydro-β-carboline carboxylic acid), previously identified in serum samples of this study (Renai, et al., 2022), xanthine pathway metabolites (i.e., dimethyl-uric acid, caffeine), and terpene derivatives (i.e., curcumenol). Regarding xanthine derivatives, the identification of uric acid derivatives is in accordance with literature information, which reports a correlation between the intake of phenolic-rich foods with the increase of uric acid levels (Lotito & Frei, 2006). However, the occurrence of caffeine has never been reported in association with berries consumption, and can be derived from caffeine-rich products consumed before fasting (Martínez-López, Sarriá, Gómez-Juaristi, Goya, Mateos, & Bravo-Clemente, 2014) and/or within the period of pool samples collection. In addition, curcumenol and its hypothesized sesquiterpene derivative were reported in this study for the first time in association with the intake of bilberry and blueberry. Finally, Gamma-CEHC, an endogenous metabolite of vitamin E already known for its anti-inflammatory properties (Wu & Croft, 2007), was identified in the molecular family, also exhibiting a positive correlation coefficient vs. time points (r = 0.441). Interestingly, this is the first report of the metabolic behaviour of this metabolite related to berry consumption.

To further extend the putative annotation of the unknown nodes in the identified molecular families, **Figure S6** of the *Supplementary materials* displays the MS/MS spectra and fragmentation schemes for the hypothesized structures (labelled by “gear” symbols) of **Fig. 4**. It is encouraging that 7 out of 8 hypothesized metabolites were characterized by a mass accuracy below 5 ppm.

As done for NI dataset, the impact of the interventions on the global urine metabolome was estimated by summing IDs spectral counts in PI mode. Differently from NI dataset, 54% of the nodes were found after the VM supplement intake, whereas the 46% were found after VC intake.

### 3.4 FBMN vs. conventional annotation workflow for metabolite discovery

In order to investigate the increased discovery capacity and annotation coverage of endogenous and exogenous urinary metabolites of the FBMN approach, a conventional selection and annotation process of urinary metabolites was performed on a statistical basis in relation to their post-prandial behaviour, as elsewhere performed in nutrimetabolomics (Renai, et al., 2022). Overall, the latter approach resulted in 50 and 106 statistically significant intra-intervention features in PI and NI datasets, respectively. However, the manual structure elucidation allowed to tentatively identify a restricted set of only 18 metabolites. Based on AUC analysis, 4 of them were found to be significant for VC, whereas 14 were significantly associated with VM. The tentatively identified metabolites by intra-intervention statistical analysis are reported in **Table S2** in section S7 of the *Supplementary materials*. Overall, the annotation coverage of the urinary metabolome was extended of the 72% by FBMN approach, in comparison with the conventional workflow for selection and annotation. Comparing findings from the statistical analyses with those obtained by the FBMN approach, hydroxyhippuric acid and dihydrocaffeic acid glucuronide were identified with both approaches, confirming their correlation with the VM and VC interventions here investigated. Moreover, postprandial statistical analysis confirmed the occurrence of groups of structurally-related metabolites that were significantly altered upon berry intake, identified also with FBMN, belonging to furoic acid derivatives (**Fig. 2B**), hydroxy and/or methoxy benzoic acids (**Figs. 2A, 3**, and **4**), and abscisic acid derivatives (**Fig. 1**). Interestingly, postprandial statistical analysis allowed to identify the metabolite categories of valerolactone and valeric acid derivatives, which are colon-derived catabolites of flavanols (e.g., procyanidins, catechin and epicatechin). These metabolites are frequently found in biological fluids after consumption of fruits and vegetables (Iglesias-Carres, Mas-Capdevila, Bravo, Aragonès, Arola-Arnal, & Muguerza, 2019) and in this study significantly characterized the VM intervention. The features related to these putatively annotated metabolites by the conventional approach, occurred also inside the FBMN networks. In detail, features at *m/z* 383.10 (dihydroxyphenyl-γ-valerolactone glucuronide II) and 287.02 (dihydroxyphenyl-γ-valerolactone sulfate) occurred inside the same molecular family inside NI dataset, sharing the common MS/MS bas peak at *m/z* 207.06 (**Figure S7A** and **S7B** of the *Supplementary materials*) corresponding to the loss of glucuronide and sulfate moieties for the former and the latter precursor ions, respectively. Inside PI dataset, only the feature at *m/z* 289.04 (dihydroxyphenyl-γ-valerolactone sulfate) gave rise to a singlet in the network, confirmed by the fragmentation pattern (**Figure S7C**) given for this node in comparison with **Table S2**.

Due to the lack of available analytical standards, and thus of library references, FBMN was not able to identify this interesting molecular families. This latter aspect highlights the importance and the future need of expanding the coverage of on-line spectra libraries, which is a fundamental step to increase the number of relevant and reliable metabolites annotations, as was demonstrated by the “In-house” library presented in this study.

## 4. Conclusions

Urine is a reservoir of transformation products resulting from co□metabolic and microbiome processing of foods and xenobiotics. Its complex composition poses difficulties in traditional metabolite annotation processes, which frequently result in a low-informative outcome.

This research investigated for the first time the use of the FBMN approach for the annotation of postprandial metabolites in comparison with a conventional identification workflow commonly adopted for discovering significant postprandial metabolites in nutrimetabolomics interventions. Accordingly, the FBMN approach was applied to a two-arms intervention study on the intake of VM and VC supplements, annotating with high mass accuracy 23 and 42 metabolites in NI and PI mode, respectively, thus increasing the annotation capacity of the 72% in comparison with a conventional selection and annotation workflow. Hence, FBMN resulted in an efficient analytical tool to extend the annotation coverage in complex metabolomics datasets. This result was obtained thanks to the optimization of MN input options in combination with the coupling of an on purpose built “In-house” spectral library with public spectral libraries. Additionally, the structures of fifteen metabolites were hypothesized by spectral analysis enabled by mass spectral networking, allowing for a deeper knowledge on the potential transformation involved in human metabolism.

Thanks to the quantitative information introduced by FBMN approach and the correlation analysis performed of metabolite abundance vs. time points, it was possible to present a first attempt for estimating the metabolic origin (i.e., phase I and phase II metabolism) of the several molecular families, some of them confirmed also with the conventional approach. Furthermore, the metabolite postprandial response in NI (VM: 38%, VC: 62%) and PI (VM: 54%, VC: 46%) datasets was highlighted for identified nodes, providing an idea of the potentially wide and differential impact of the two bilberry intakes on the global urine metabolome.

Taken together, the proposed approach will not only help to confirm current knowledge of phenolic-rich food metabolism, but will also allow to expand our understanding of the complex urinary metabolome by annotating additional food-derived as well as endogenous metabolites.

## Supporting information

List of Referece Standards used to build the library

Metadata and Library Information

## Declaration of interest

The authors declare that they have no conflict of interest.

## Acknowledgements

The authors wish to acknowledge the support of the Regione Toscana and the private companies Il Baggiolo S.r.l., Danti Giampiero & C. S.n.c., Azienda Agricola Il Sottobosco, and Farmaceutica MEV S.r.l., through the PRAF Misura 1.2. e) grant.

## Author contributions

Lapo Renai: conceptualization, investigation, software, formal analysis, data curation, writing - original draft; Marynka Ulaszewska: conceptualization, software, data curation, sample analysis and annotation, formal analysis, supervision, writing - review & editing; Fulvio Mattivi: coordinating metabolomics experiments, project administration; Riccardo Bartoletti: clinical trial execution, project administration; Massimo Del Bubba: project administration, funding acquisition, writing - review & editing; Justin J. J. van der Hooft: conceptualization, formal analysis, writing - review & editing, supervision.

## Notes

### Competing Interest Statement

The authors have declared no competing interest.

## References

Ancillotti, C., Ciofi, L., Rossini, D., Chiuminatto, U., Stahl-Zeng, J., Orlandini, S., Furlanetto, S., & Del Bubba, M. (2017). Liquid chromatographic/electrospray ionization quadrupole/time of flight tandem mass spectrometric study of polyphenolic composition of different Vaccinium berry species and their comparative evaluation. Analytical and bioanalytical chemistry, 409(5), 1347–1368. DOI 10.1007/s00216-016-0067-y

Ancillotti, C., Ulaszewska, M., Mattivi, F., & Del Bubba, M. (2019). Untargeted metabolomics analytical strategy based on liquid chromatography/electrospray ionization linear ion trap quadrupole/orbitrap mass spectrometry for discovering new polyphenol metabolites in human biofluids after acute ingestion of vaccinium myrtillus berry supplement. Journal of the American Society for Mass Spectrometry, 30(3), 381–402. https://doi.org/10.1007/s13361-018-2111-y

Aravind, S. M., Wichienchot, S., Tsao, R., Ramakrishnan, S., & Chakkaravarthi, S. (2021). Role of dietary polyphenols on gut microbiota, their metabolites and health benefits. Food Research International, 110189. https://doi.org/10.1016/j.foodres.2021.110189

Aron, A. T., Gentry, E. C., McPhail, K. L., Nothias, L.-F., Nothias-Esposito, M., Bouslimani, A., Petras, D., Gauglitz, J. M., Sikora, N., & Vargas, F. (2020). Reproducible molecular networking of untargeted mass spectrometry data using GNPS. Nature protocols, 15(6), 1954–1991. https://econewstoday.com/

Beniddir, M. A., Kang, K. B., Genta-Jouve, G., Huber, F., Rogers, S., & van der Hooft, J. J. (2021). Advances in decomposing complex metabolite mixtures using substructure-and network-based computational metabolomics approaches. Natural product reports. https://doi.org/10.1039/D1NP00023C

Chaleckis, R., Meister, I., Zhang, P., & Wheelock, C. E. (2019). Challenges, progress and promises of metabolite annotation for LC–MS-based metabolomics. Current opinion in biotechnology, 55, 44–50. https://doi.org/10.1016/j.copbio.2018.07.010

Colak, N., Primetta, A. K., Riihinen, K. R., Jaakola, L., Grúz, J., Strnad, M., Torun, H., & Ayaz, F. (2017). Phenolic compounds and antioxidant capacity in different-colored and non-pigmented berries of bilberry (Vaccinium myrtillus L.). Food Bioscience, 20, 67–78. https://doi.org/10.1016/j.fbio.2017.06.004

de Mello, V. D., Lankinen, M. A., Lindström, J., Puupponen□Pimiä, R., Laaksonen, D. E., Pihlajamäki, J., Lehtonen, M., Uusitupa, M., Tuomilehto, J., & Kolehmainen, M. (2017). Fasting serum hippuric acid is elevated after bilberry (Vaccinium myrtillus) consumption and associates with improvement of fasting glucose levels and insulin secretion in persons at high risk of developing type 2 diabetes. Molecular nutrition & food research, 61(9), 1700019. https://doi.org/10.1002/mnfr.201700019

Degu, A., Ayenew, B., Cramer, G. R., & Fait, A. (2016). Polyphenolic responses of grapevine berries to light, temperature, oxidative stress, abscisic acid and jasmonic acid show specific developmental-dependent degrees of metabolic resilience to perturbation. Food chemistry, 212, 828–836. https://doi.org/10.1016/j.foodchem.2016.05.164

Du, X., Finn, C. E., & Qian, M. C. (2010). Volatile composition and odour-activity value of thornless ‘Black Diamond’and ‘Marion’blackberries. Food chemistry, 119(3), 1127–1134. https://doi.org/10.1016/j.foodchem.2009.08.024

Elsharif, S. A., & Buettner, A. (2016). Structure–odor relationship study on geraniol, nerol, and their synthesized oxygenated derivatives. Journal of Agricultural and Food Chemistry, 66(10), 2324–2333. https://doi.org/10.1021/acs.jafc.6b04534

Garcia□Aloy, M., Ulaszewska, M., Franceschi, P., Estruel□Amades, S., Weinert, C. H., Tor□Roca, A., Urpi□Sarda, M., Mattivi, F., & Andres□Lacueva, C. (2020). Discovery of intake biomarkers of lentils, chickpeas, and white beans by untargeted LC–MS metabolomics in serum and urine. Molecular nutrition & food research, 64(13), 1901137. https://doi.org/10.1002/mnfr.201901137

Gutsche, B., Grun, C., Scheutzow, D., & Herderich, M. (1999). Tryptophan glycoconjugates in food and human urine. Biochemical journal, 343(1), 11–19. https://doi.org/10.1042/bj3430011

Iglesias-Carres, L., Mas-Capdevila, A., Bravo, F. I., Aragonès, G., Arola-Arnal, A., & Muguerza, (2019). A comparative study on the bioavailability of phenolic compounds from organic and nonorganic red grapes. Food chemistry, 299, 125092. https://doi.org/10.1016/j.foodchem.2019.125092

Kind, T., & Fiehn, O. (2007). Seven Golden Rules for heuristic filtering of molecular formulas obtained by accurate mass spectrometry. BMC bioinformatics, 8(1), 1–20. https://doi.org/10.1186/1471-2105-8-105

Lotito, S. B., & Frei, B. (2006). Consumption of flavonoid-rich foods and increased plasma antioxidant capacity in humans: cause, consequence, or epiphenomenon? Free Radical Biology and Medicine, 41(12), 1727–1746. https://doi.org/10.1016/j.freeradbiomed.2006.04.033

Martínez-López, S., Sarriá, B., Gómez-Juaristi, M., Goya, L., Mateos, R., & Bravo-Clemente, L. (2014). Theobromine, caffeine, and theophylline metabolites in human plasma and urine after consumption of soluble cocoa products with different methylxanthine contents. Food Research International, 63, 446–455. https://doi.org/10.1016/j.foodres.2014.03.009

Medjakovic, S., & Jungbauer, A. (2013). Pomegranate: a fruit that ameliorates metabolic syndrome. Food & function, 4(1), 19–39. https://doi.org/10.1039/C2FO30034F

Neto, F. C., & Raftery, D. (2021). Expanding Urinary Metabolite Annotation through Integrated Mass Spectral Similarity Networking. Analytical chemistry. https://doi.org/10.1021/acs.analchem.1c02041

Nothias, L.-F., Petras, D., Schmid, R., Dührkop, K., Rainer, J., Sarvepalli, A., Protsyuk, I., Ernst, M., Tsugawa, H., & Fleischauer, M. (2020). Feature-based molecular networking in the GNPS analysis environment. Nature methods, 17(9), 905–908. https://doi.org/10.1038/s41592-020-0933-6

Oberleitner, D., Schmid, R., Schulz, W., Bergmann, A., & Achten, C. (2021). Feature-based molecular networking for identification of organic micropollutants including metabolites by non-target analysis applied to riverbank filtration. Analytical and bioanalytical chemistry, 1–10. https://doi.org/10.1007/s00216-021-03500-7

Padilla-González, G. F., Sadgrove, N. J., Ccana-Ccapatinta, G. V., Leuner, O., & Fernandez-Cusimamani, E. (2020). Feature-based molecular networking to target the isolation of new caffeic acid esters from yacon (Smallanthus sonchifolius, Asteraceae). Metabolites, 10(10), 407. https://doi.org/10.3390/metabo10100407

Pluskal, T., Castillo, S., Villar-Briones, A., & Orešič, M. (2010). MZmine 2: modular framework for processing, visualizing, and analyzing mass spectrometry-based molecular profile data. BMC bioinformatics, 11(1), 1–11. https://doi.org/10.1186/1471-2105-11-395

Quinn, R. A., Melnik, A. V., Vrbanac, A., Fu, T., Patras, K. A., Christy, M. P., Bodai, Z., Belda-Ferre, P., Tripathi, A., & Chung, L. K. (2020). Global chemical effects of the microbiome include new bile-acid conjugations. Nature, 579(7797), 123–129. https://doi.org/10.1038/s41586-020-2047-9

Ramos, A. E. F., Evanno, L., Poupon, E., Champy, P., & Beniddir, M. A. (2019). Natural products targeting strategies involving molecular networking: Different manners, one goal. Natural product reports, 36(7), 960–980. https://doi.org/10.1039/C9NP00006B

Renai, L., Ancillotti, C., Ulaszewska, M., Garcia-Aloy, M., Mattivi, F., Bartoletti, R., & Del Bubba, M. (2022). Comparison of chemometrics strategies for potential exposure markers discovery and false positive reduction in untargeted metabolomics: application to the serum analysis by LC-HRMS after intake of Vaccinium fruits supplements. Analitycal and bioanalytical chemistry, in press.

Rivera-Mondragón, A., Tuenter, E., Ortiz, O., Sakavitsi, M. E., Nikou, T., Halabalaki, M., Caballero-George, C., Apers, S., Pieters, L., & Foubert, K. (2020). UPLC-MS/MS-based molecular networking and NMR structural determination for the untargeted phytochemical characterization of the fruit of Crescentia cujete (Bignoniaceae). Phytochemistry, 177, 112438.

Said, I. H., Truex, J. D., Heidorn, C., Retta, M. B., Petrov, D. D., Haka, S., & Kuhnert, N. (2020). LC-MS/MS based molecular networking approach for the identification of cocoa phenolic metabolites in human urine. Food Research International, 132, 109119. https://doi.org/10.1016/j.foodres.2020.109119

Schrimpe-Rutledge, A. C., Codreanu, S. G., Sherrod, S. D., & McLean, J. A. (2016). Untargeted metabolomics strategies—challenges and emerging directions. Journal of the American Society for Mass Spectrometry, 27(12), 1897–1905. https://doi.org/10.1007/s13361-016-1469-y

Shannon, P., Markiel, A., Ozier, O., Baliga, N. S., Wang, J. T., Ramage, D., Amin, N., Schwikowski, B., & Ideker, T. (2003). Cytoscape: a software environment for integrated models of biomolecular interaction networks. Genome research, 13(11), 2498–2504. doi: 10.1101/gr.1239303

Spicer, R. A., Salek, R., & Steinbeck, C. (2017a). Compliance with minimum information guidelines in public metabolomics repositories. Scientific data, 4(1), 1–8. https://doi.org/10.1038/sdata.2017.137

Spicer, R. A., Salek, R., & Steinbeck, C. (2017b). A decade after the metabolomics standards initiative it’s time for a revision. Scientific data, 4(1), 1–3. https://doi.org/10.1038/sdata.2017.138

Stevens, J. F., & Maier, C. S. (2016). The chemistry of gut microbial metabolism of polyphenols. Phytochemistry Reviews, 15(3), 425–444. DOI 10.1007/s11101-016-9459-z

Sumner, L. W., Amberg, A., Barrett, D., Beale, M. H., Beger, R., Daykin, C. A., Fan, T. W.-M., Fiehn, O., Goodacre, R., & Griffin, J. L. (2007). Proposed minimum reporting standards for chemical analysis. Metabolomics, 3(3), 211–221. https://doi.org/10.1007/s11306-007-0082-

Van Der Hooft, J. J., Padmanabhan, S., Burgess, K. E., & Barrett, M. P. (2016). Urinary antihypertensive drug metabolite screening using molecular networking coupled to high-resolution mass spectrometry fragmentation. Metabolomics, 12(7), 1–15. https://doi.org/10.1007/s11306-016-1064-z

Vargas, F., Weldon, K. C., Sikora, N., Wang, M., Zhang, Z., Gentry, E. C., Panitchpakdi, M. W., Caraballo□Rodríguez, A. M., Dorrestein, P. C., & Jarmusch, A. K. (2020). Protocol for community□created public MS/MS reference spectra within the Global Natural Products Social Molecular Networking infrastructure. Rapid Communications in Mass Spectrometry, 34(10), e8725. https://doi.org/10.1002/rcm.8725

Vetrani, C., Rivellese, A. A., Annuzzi, G., Adiels, M., Borén, J., Mattila, I., Orešič, M., & Aura, A.-M. (2016). Metabolic transformations of dietary polyphenols: comparison between in vitro colonic and hepatic models and in vivo urinary metabolites. The Journal of nutritional biochemistry, 33, 111–118. https://doi.org/10.1016/j.jnutbio.2016.03.007

Wu, J., & Croft, K. (2007). Vitamin E metabolism. Molecular aspects of medicine, 28(5-6), 437–452. https://doi.org/10.1016/j.mam.2006.12.007

Xie, H.-F., Kong, Y.-S., Li, R.-Z., Nothias, L.-F., Melnik, A. V., Zhang, H., Liu, L.-L., An, T.-T., Liu, R., & Yang, Z. (2020). Feature-Based Molecular Networking Analysis of the Metabolites Produced by in vitro Solid-State Fermentation Reveals Pathways for the Bioconversion of Epigallocatechin Gallate. Journal of Agricultural and Food Chemistry, 68(30), 7995–8007. https://doi.org/10.1021/acs.jafc.0c02983

Zhang, H., Ma, Z. F., Luo, X., & Li, X. (2018). Effects of mulberry fruit (Morus alba L.) consumption on health outcomes: A mini-review. Antioxidants, 7(5), 69. https://doi.org/10.3390/antiox7050069

