## Supplementary materials for "Boosting annotation in nutrimetabolomics by Feature-Based Molecular Networking: analytical and computational strategies applied to human urine samples from an untargeted LC-MS/MS based bilberry-blueberry intervention study"

### S1. Chemicals and reagents

Methanol and acetonitrile LC-MS Ultra CHROMASOLV™ and formic acid were purchased from Sigma Aldrich (St. Louis, MO, USA). The ultrapure water was obtained by purifying demineralized water in a Milli-Q system from Millipore (Bedford, MA, USA). The internal standard trans-cinnamic-d5 acid (CIN-d5) was purchased from CDN ISOTOPES Inc. (Pointe-Claire, Quebec, Canada). The surrogate standards L-tryptophan- 2′,4′,5′,6′,7′-d5 (TRY-d5) and N-benzoyl-d5-glycine (hippuric acid-d5, HIP-d5) were obtained from Sigma Aldrich and CDN ISOTOPES Inc., respectively.

### S2. Supplement characterization

In order to assess the amount of total soluble polyphenols (TSP), total monomeric anthocyanins (TMA) and the most important phenolic compounds commonly found in bilberries and blueberries, the following protocols were adopted. About 500 mg aliquots of supplement were homogenized in an ice bath under magnetic stirring with 15 mL of a methanol/water solution 8/2 (v/v) containing 10 mM NaF to inactivate polyphenol oxidase; the mixture was centrifuged at 1800xg for 5 min and the supernatant recovered. This procedure was repeated twice and the extracts combined and analysed for TSP, TMA, and selected phenolic compounds. TSP were spectrophotometrically determined as following described. Extract aliquots (100-200 μL, depending on the TSP concentration in the extract) were mixed with 200 μL of Folin-Ciocalteau reagent. After 3 min, 400 μL of an aqueous solution saturated with sodium carbonate were added and the mixture obtained was made up to 10 mL with ultra-pure water. The solution was dark incubated for 1 h. Afterwards, the absorbance was measured at 740 nm and TSP concentration calculated on the basis of a catechin calibration curve; accordingly, the results were expressed as milligrams of catechin/g of supplement. TMA were determined with the pH differential method using cyanidin-3-glucoside as reference standard. Aliquots of 100-200 µL of extract were diluted in buffer solutions at pH=1 and pH=4.5, so as to obtain a final volume of 10 mL. The absorbance (Abs) of both solutions were measured at 520 and 700 nm and the quantity “ΔAbs” was calculated according to equation 1.

$\Delta Abs=\left( \mathrm{Abs}_{pH=1}^{520 nm}-\mathrm{Abs}_{pH=1}^{700 nm} \right)-\left( \mathrm{Abs}_{pH=4.5}^{520 nm}-\mathrm{Abs}_{pH=4.5}^{700 nm} \right)$	(1)

Similarly, “ΔAbs” values were also calculated for different concentrations of cyanidin-3-glucoside reference standard and plotted as a function of corresponding cyanidin-3-glucoside concentrations. The equation best fitting the experimental data was calculated by the least square regression method, thus obtaining a linear calibration curve, which was used for measuring TMA in the extracts. The results were expressed as milligrams of cyanidin-3-glucoside per g of supplement.

**Table S1 –** Total soluble polyphenol (TSP) and total monomeric anthocyanin (TMA) concentrations of *V. myrtillus* and *V. corymbosum* supplements administered to volunteers during the study.

|  | *V. myrtillus* supplement | *V. corymbosum* supplement |
| --- | --- | --- |
| TSP (mg CAT/g supplement dry weight) | 38.8 ± 2.1 | 9.3 ± 1.0 |
| TMA (mg CYA-3-GLU/g supplement dry weight) | 30.1 ± 2.8 | 6.4 ± 0.9 |

### S3.  Sample extraction

100 µL of urine were placed into the Millipore 96-well plate (Waters, Milford, MA, USA)and 100 µL of surrogate standards in methanol (25 µg/mL TRI-d5 and HIP-d5) were added. Then, a cap mat was applied on the well plate and the system was mixed for 5 minutes. After mixing, the well plate and the collection plate were placed on the positive pressure processor (Positive Pressure-96 Processor, Waters) and the filtration was performed setting the flow for 60 psi for 5 minutes. Then, 300 µL of internal standard (0.83 µg/mL CIN-d5) were added to each position of the collection plate and the system was mixed for 5 minutes. The resulting filtrated and diluted urine samples were transferred in labelled brown vials for LC-MS analysis. For testing the urine filtration procedure, a quality control (QC) sample consisting of a pool of the same volume of all urine samples was prepared. The same extraction protocol was performed on QC aliquots randomly placed in the well plate. The QC extracts together with the extraction blanks were injected before the LC-MS analysis of individual samples.

### S4. LC-MS/MS analysis

The chromatographic analysis was performed on a Kinetex C18 column and a guard column containing the same stationary phase. Column temperature was set at 40°C. Water (eluent A) and acetonitrile (eluent B), both with 0.1 % formic acid, were used as mobile phases. For chromatographic analysis of urine sample, the following gradient was adopted: 0-1 min isocratic 5% B, 1-7 min linear gradient 5-45% B, 7-8.5 min linear gradient 45-80% B, 8.5-10.5 min isocratic 80% B, 10.5 -11 min linear gradient 80-5% B, 11-12 min 5% B. For chromatographic analysis of serum samples, the following gradient was adopted: 0-1 min isocratic 5% B, 1-12.5 min linear gradient 5-100 % B, 12.5-14 min isocratic 100% B, 14-14.3 min linear gradient 100-5% B, 14.3-15.3 min 5% B. For both urine and serum analysis, the flow rate was 350 µL/min and the injection volume was 5 μL. The ESI conditions in positive (and negative) mode were: spray voltage 5.0 kV (-3.5 kV), heated capillary temperature 320ºC, capillary voltage 35 V (-35 V) and tube lens 110 V (-110 V). In the LTQ component of the instrument, nitrogen was used as both the sheath gas (35 U) and auxiliary gas (5 U), and helium was used as the damping gas. All measurements were done using the automatic gain control of the LTQ to adjust the number of ions entering the trap. Mass calibration was performed with every sequence run just prior to starting the batch by using flow injection of the manufacturer’s calibration standards mixture allowing for mass accuracies lower than 5 ppm in external calibration mode. Data-dependent acquisition was performed to obtain both full scan (mass range from 100 to 1000 Da) and MS/MS spectra with a resolving power for MS/MS scans of 7500. Product ions were generated in the LTQ trap at collision energy 35 eV using an isolation width of 2 Da.

### S5. Molecular networking analyses

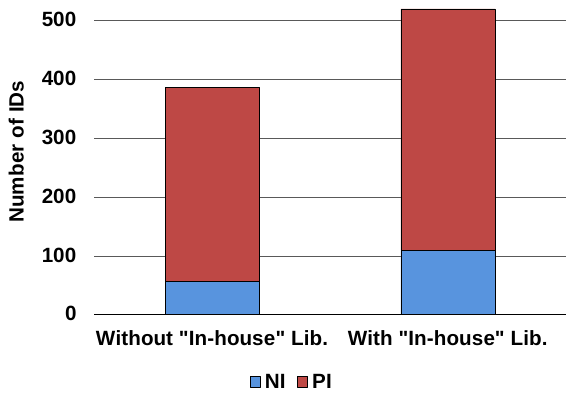

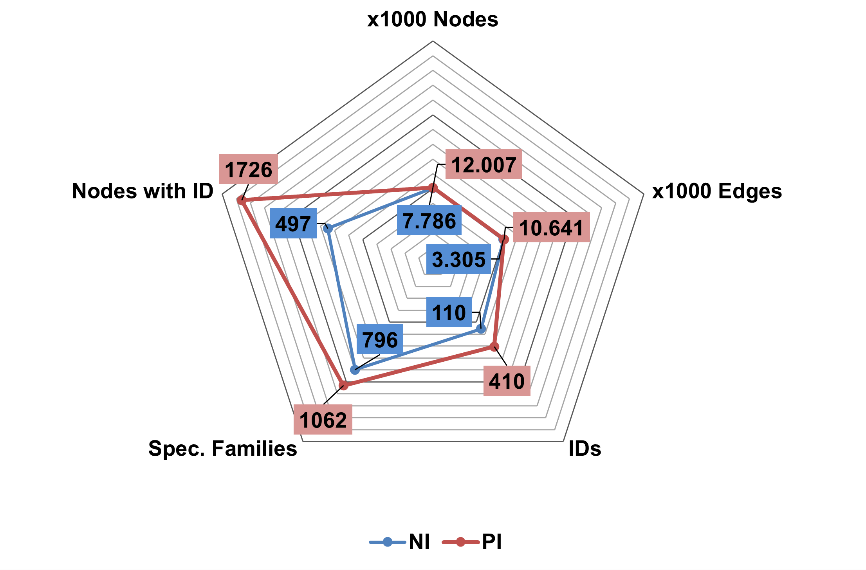

**(B)**

**(A)**

**(A)**

**Figure S1** − (A) Number of identified nodes (IDs found by Molecular Networking) from both negative ionisation and positive ionisation datasets before and after the inclusion of the “In-house” library. (B) Radar graph of networking outputs obtained with the selected basic and advanced options (see paragraph 2.7 of the main text) selected for Feature-Based Molecular Networking workflow for both negative ionisation (blue line) and positive ionisation (red line) datasets.

### S6. MS/MS Spectra and schemes for hypothesized structures

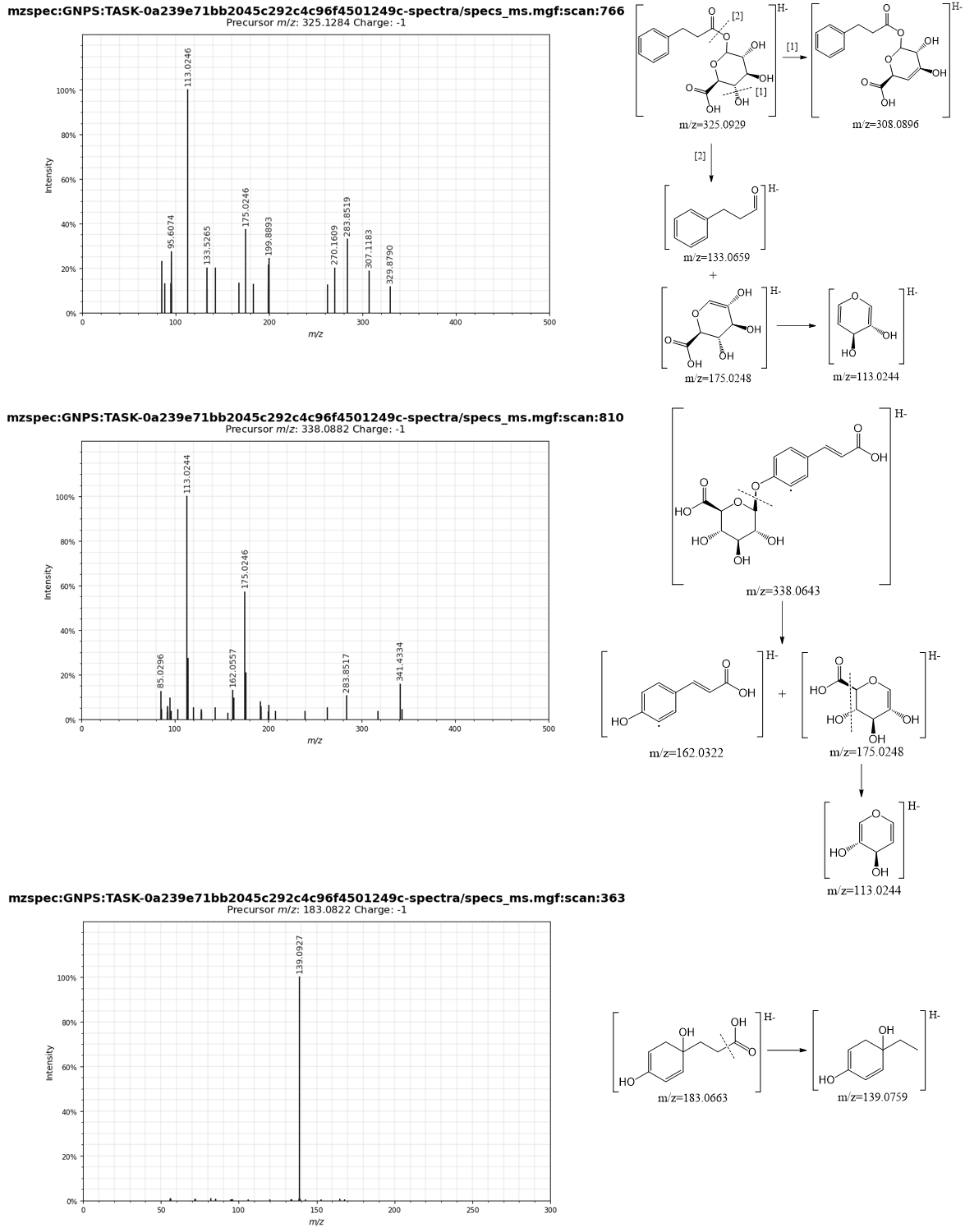

**Figure S2** – MS/MS spectra and hypothesized fragmentation scheme of putative metabolites reconstructed from network analysis of Figure 1 of the main text.

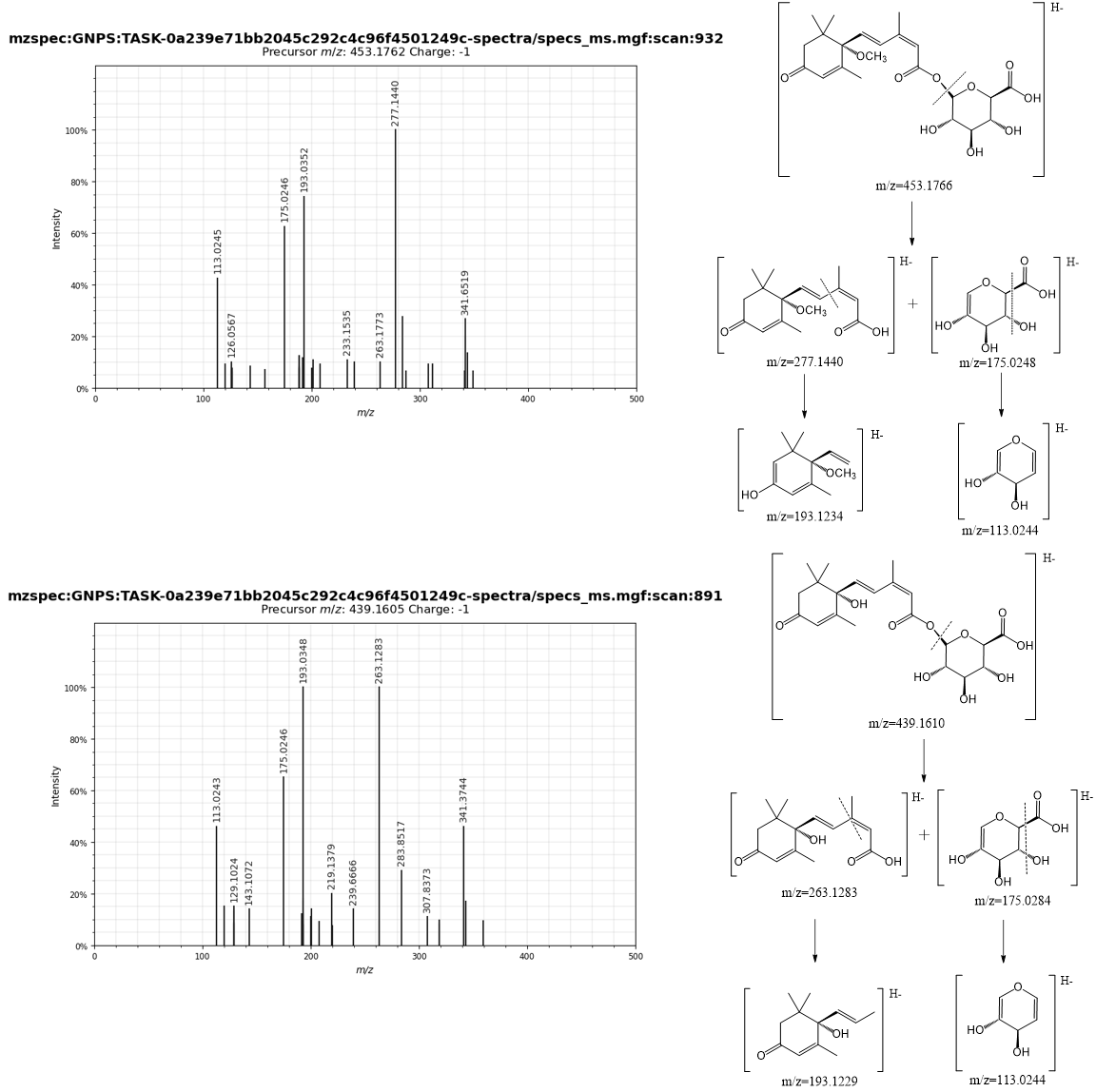

**Figure S3** – MS/MS spectra and hypothesized fragmentation scheme of putative metabolites methoxyabscisic acid glucuronide (top) and abscisic acid glucuronide (bottom), reconstructed from network analysis of Figure 1 of the main text.

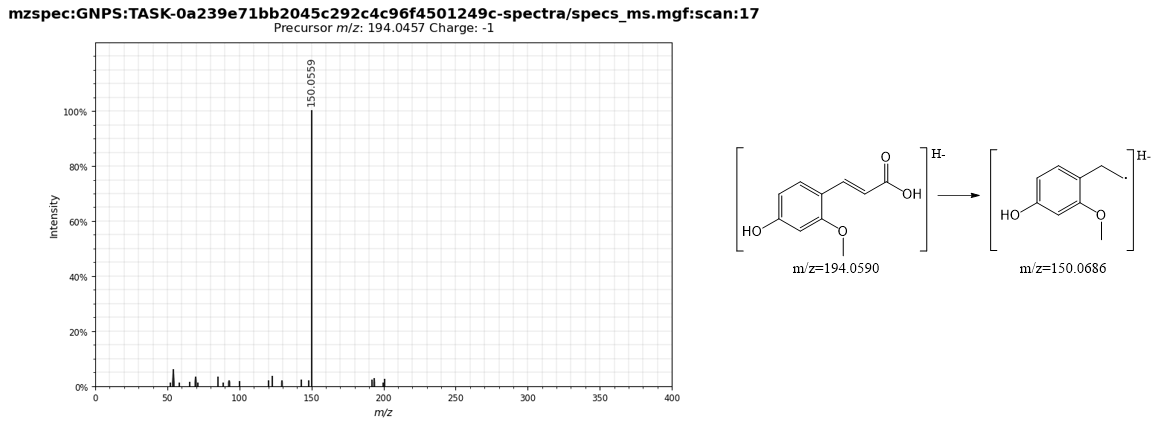

**Figure S4** – MS/MS spectra and hypothesized fragmentation scheme of putative metabolite reconstructed from network analysis of Figure 2B of the main text.

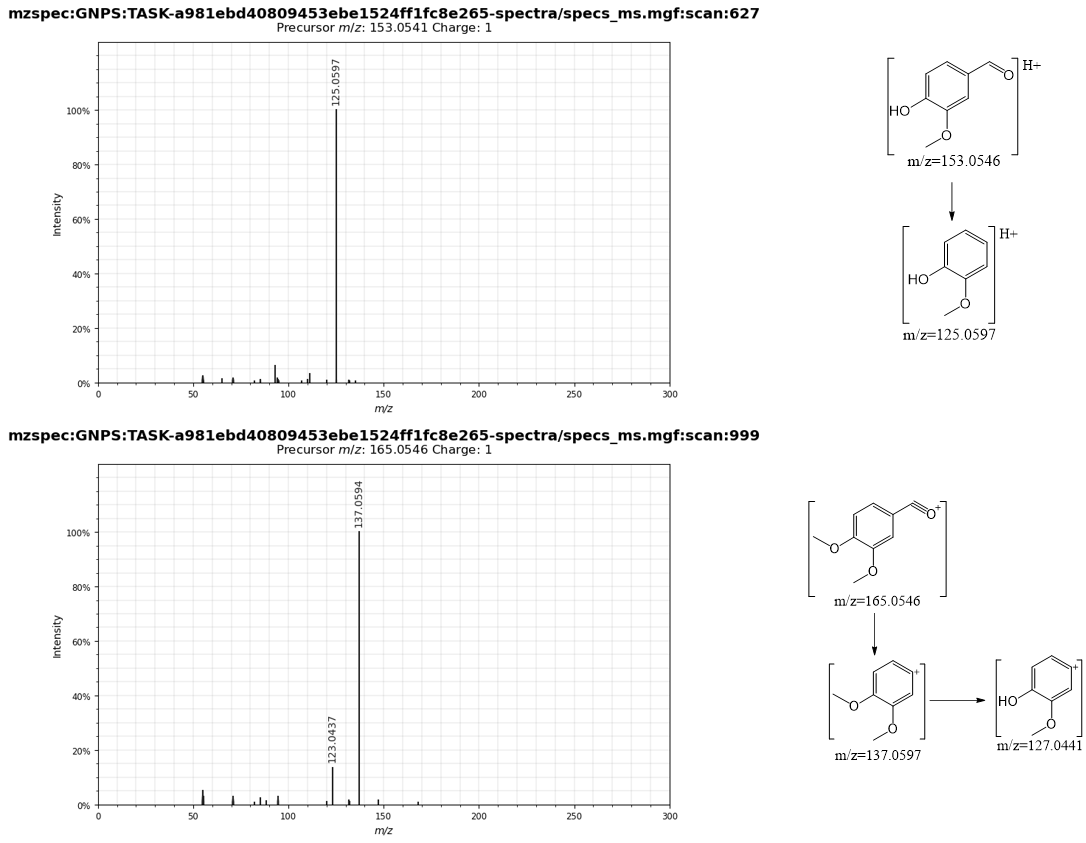

**Figure S5** – MS/MS spectra and hypothesized fragmentation scheme of putative metabolites reconstructed from network analysis of Figure 4 of the main text.

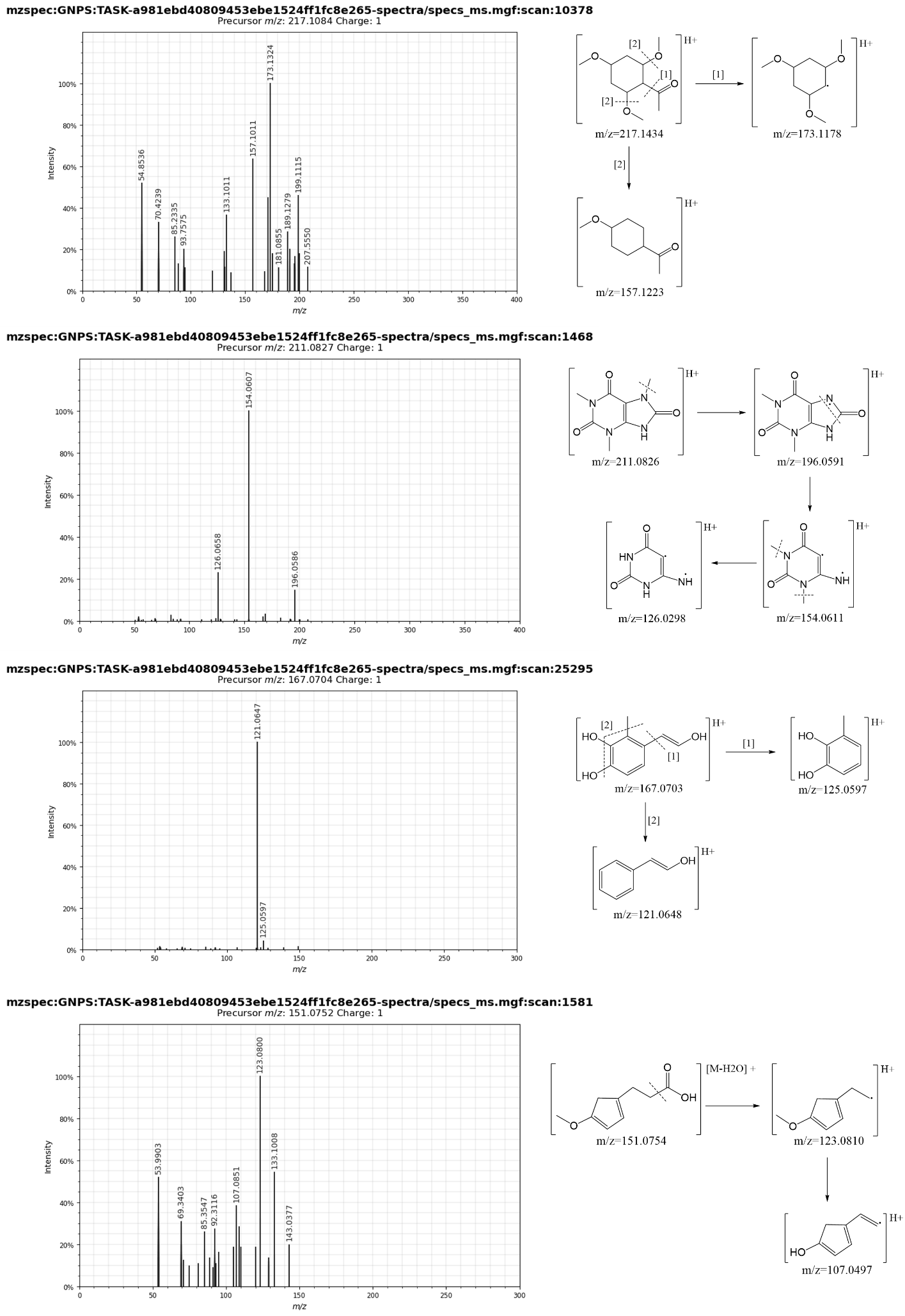

**Figure S6** – MS/MS spectra and hypothesized fragmentation scheme of putative metabolites reconstructed from network analysis of Figure 5 of the main text.

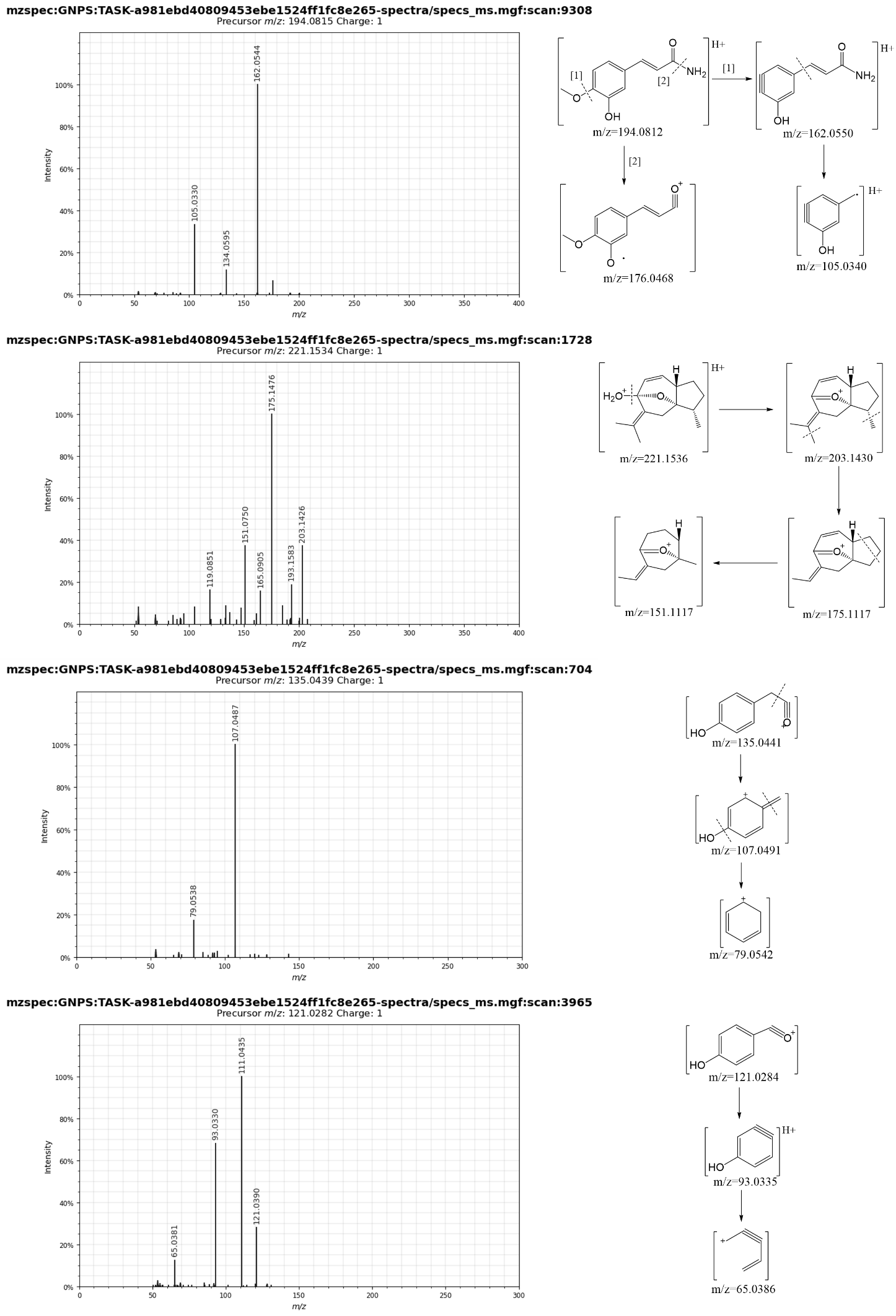

**Figure S6** – Continued.

### S7. FBMN vs. conventional annotation workflow for metabolite discovery

**Table S2** – List of metabolites tentatively identified by traditional statistical analysis in urine samples after acute ingestion of VM and VC supplements. For each statistically significant feature we reported retention time (t_R_), precursor ion exact mass (Da) and charge state in squared brackets, product ions (MS/MS, Da), the tentative identification and the postprandial statistical significance.

| t_R_ | Precursor ion | MS/MS | Tentative identification | *P*-Value & diet |
| --- | --- | --- | --- | --- |
| 0.96 | 128.0191 [M+H]+ | MS^2^ 128: 110.0085 (100); MS^3^ 128-110: 82.0134 (100). | Hydroxy-furoic acid | 8.55E-03, VC |
| 1.42 | 369.0514 [M+H]+ | MS^2^ 369: 193.0968 (100), 175.0870 (15). | dihydroxytryptamine/oxoamide glucuronide | 1.01E-02, VM |
| 2.24 | 220.1176 [M+H]+ | MS^2^ 220: 202.1066 (100), 184.0962 (40), 174.1117(10), 90.0544 (40). | Pantothenic acid | 2.01E-02, VM |
| 3.07 | 194.0451 [M+H]+ | MS^2^ 194: 150.0554 (100), 194.0447 (60), 93.0344(17), 148.0399 (12). | Hydroxyhippuric acid | 5.24E-03, VM |
|  | 389.0967 [2M-H]- | MS^2^ 389: 194.0447 (100), 150.0555 (12). |  |  |
| 3.33 | 373.0769 [M-H]- | MS^2^ 373: 197.0452 (100), 182.0212 (15), 175.0247(40), 113.0243 (30). | Syringic acid glucuronide I | 5.02E-04, VC |
| 3.54 | 505.1190 [M-H]- | MS^2^ 505: 343.0662 (80), 329.0871(100), 191.0346(60), 167.0348(60); MS^3^ 505-329: 123.050(100). | Vanilic acid glucoside-glucuronide | 2.60E-03, VM |
| 3.95 | 373.0769 [M-H]- | MS^2^ 373: 197.0452 (100), 182.0217(10), 175.0244 (30), 113.0244 (35). | Syringic acid glucuronide II | 2.60E-03, VC |
| 4.22 | 307.0490 [M+H]+ | MS^2^ 307: 289.0380 (100), 173.0569 (40); MS^3^ 307-289: 229.0160 (100), 209.0803 (70), 191.0696 (60) | Dihydroxyphenyl-valeric acid sulfate | 8.55E-03, VM |
| 4.27 | 449.1080 [M+H]+ | MS^2^ 449: 287.0543 (100), 317.0655 (30), 143.0432 (5). | Cyanidin-hexoside | 2.00E-03, VM |
| 4.54 | 339.07139 [M-H]- | MS^2^ 339: 175.0248 (100), 163.0394 (40). | Hydroxycoumarin glucuronide | 1.56E-02, VM |
| 4.60 | 385.1120 [M+H]+ | MS^2^ 209: 191.0697 (100), 149.0593 (44), 163.0755 (5), 123.0442 (4). | Dihydroxyphenyl-γ-valerolactone glucuronide I | 2.93E-03, VM |
| 4.80 | 383.0975 [M-H]- | MS^2^ 383: 207.0658 (100), 175.0244 (40), 113.0243 (45), 163.0767 (20). | Dihydroxyphenyl-γ-valerolactone glucuronide II | 3.96E-03, VM |
| 4.97 | 289.0373 [M+H]+ | MS^2^ 289: 271.0267 (35), 229.0163 (100), 209.0805(50), 191.0701(55), 149.0596(30), 131.0489(60), 143.0316(15) | Dihydroxyphenyl-γ-valerolactone sulfate | 2.93E-03, VM |
|  | 287.0223 [M-H]- | MS^2^ 287: 207.0622 (100), 163.0762 (3); MS^3^ 287-207: 163.0760(100), 122.0376 (20), 109.0292 (12). |  |  |
| 5.16 | 343.0663 [M-H]- | MS^2^ 163: 175.0246 (100), 167.0349 (20), 113.0245 (30). | Vanillic acid glucuronide | 1.47E-02, VM |
| 5.18 | 357.0819 [M-H]- | MS^2^ 357: 181.0502 (100), 137.0611 (40). | Dihydrocaffeic acid glcucuronide | 7.95E-03, VM |
| 5.25 | 383.0434 [M-H]- | MS^2^ 383: 303.0858 (100), 216.9803 (59), 137.0239 (44), 245.0123 (15), 259.0965 (7). | (Epi)catechin-methyl sulfate | 1.22E-02, VM |
| 5.33 | 455.15487 [M-H]- | MS^2^ 455: 279.1232 (100), 217.1228 (30), 143.1070 (15), 175.0241 (6). | Hydroxy-abscisic acid glucuronide | 1.32E-03, VC |
| 6.11 | 333.0607[M-H]- | MS^2^ 333: 165.0191 (100), 183.0296(22), 137.0244(20), 289.0709 (19). | Methyl-dihydromyricetin | 2.60E-03, VM |

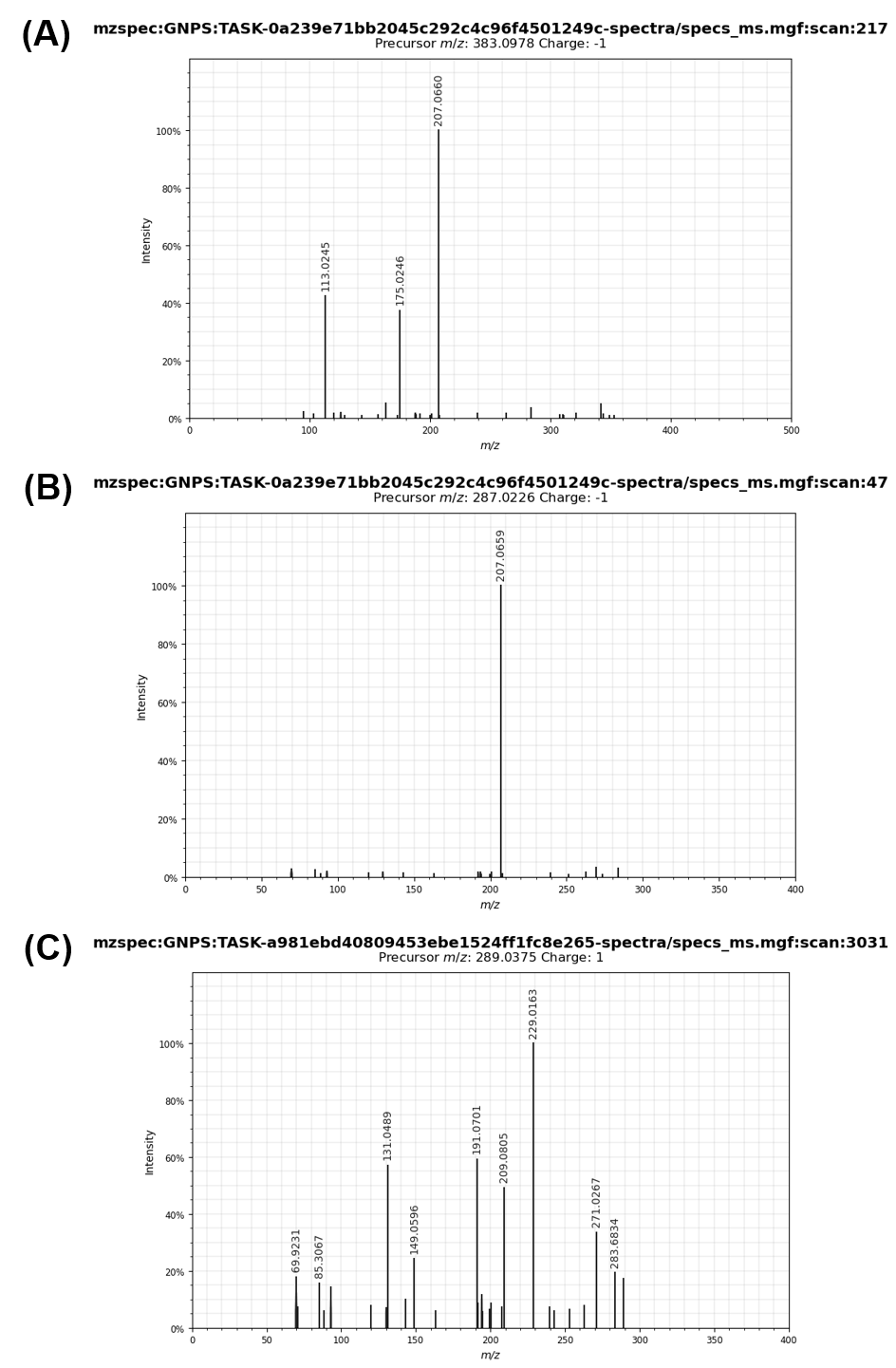

**Figure S7** – MS/MS spectra of the putative annotated valerolactone derivatives reported in Table S2 and occurring inside NI and PI FBMN. (A) Dihydroxyphenyl-γ-valerolactone glucuronide II, *m/z* 383.10 Da. (B) Dihydroxyphenyl-γ-valerolactone sulfate, *m/z* 287.02 Da. (C) Dihydroxyphenyl-γ-valerolactone sulfate, *m/z* 289.04 Da.
